# National Wetland Map 5 – An improved spatial extent and representation of inland aquatic and estuarine ecosystems in South Africa

**DOI:** 10.1101/640441

**Authors:** Heidi van Deventer, Lara van Niekerk, Janine Adams, Millicent Ketelo Dinala, Ridhwannah Gangat, Stephen J Lamberth, Mervyn Lötter, Namhla Mbona, Fiona MacKay, Jeanne L Nel, Carla-Louise Ramjukadh, Andrew Skowno, Steven P Weerts

## Abstract

The improved representation of inland aquatic (freshwater) and estuarine ecosystems and associated data was a key component of the 2018 National Biodiversity Assessment, and is an essential step in enhancing defensible land use planning and decision making. This paper reports on this enhancement of the National Wetland Map version 5 (NWM5) for South Africa and other data layers associated with the South African Inventory of Inland Aquatic Ecosystems. Detail is provided on (i) the extent of wetlands mapped in NWM5, compared to previous versions of the NWMs; (ii) the improved extent of inland wetlands mapped in focus areas in NWM5 relative to NWM4; (iii) the type of cover associated with the wetlands (inundated, vegetated or arid); (iv) the ecotone between rivers or inland wetlands and estuaries mapped as freshwater—estuarine transition zones; and (v) level of confidence for the inland wetlands in terms of how well the extent and hydrogeomorphic units were captured for each sub-quaternary catchment of South Africa. A total of 4 698 824 ha (3.9% of South Africa) of inland aquatic ecosystems and artificial wetlands have now been mapped, with NWM5 delineating 123% more inland wetlands (2 635 809 ha or 2.2% of SA) compared with NWM4. The estuarine functional zone, which encapsulates all estuarine processes, associated habitats and biota, was refined for 292 systems totalling 188 944 ha, with the addition of 42 micro-estuaries totalling 246 ha. Nearly 600 000 ha (0.5% of SA) of artificial wetlands were mapped in SA. Inland wetlands are predominantly palustrine (55%), some arid (34%) and few inundated systems (11%). Transition zones between freshwater ecosystems and estuaries formed a small fraction (<1.5%) of river total extent (164 018 km), indicating an ecotone where biota and processes continuously vary from freshwater to estuarine. The majority of inland wetlands (~70%) had a low confidence ranking for designation of extent and typing. Future improvements of the map should be focussed on catchment-based improvements particularly in strategic water-source areas, areas of high development pressure and those with low confidence designation of wetland type.

## Introduction

The South African National Wetland Map (NWM) portrays the spatial extent and ecosystem types of two of the three broad aquatic ecosystems, namely estuarine and inland aquatic (freshwater) ecosystems. An aquatic ecosystem is defined as ‘an ecosystem that is permanently or periodically inundated by flowing or standing water, or which has soils that are permanently or periodically saturated within 0.5 m of the soil surface’ (Ollis et al., 2013:1). In South Africa, inland aquatic ecosystems comprise both rivers and inland wetlands, and are distinguished from estuarine systems, although an ecotone of transition exits amongst these systems where biotic and abiotic processes vary through hydrological cycles. The NWM represents the extent and ecosystem types of the estuarine and inland wetlands, collectively known as wetlands, and informs decision makers in assessing development applications, land use and conservation planning and policy making (Nel et al., 2016). Wetlands are protected under the National Water Act (*Act 36 of 1998*; RSA, 1998a), whereas estuaries receive additional protection under the Marine Living Resources Act (*Act No. 18 of 1998*; RSA 1998b) and National Environmental Management: Integrated Coastal Management Act (*Act No. 24 of 2008*; RSA 2008). The spatial representation of the aquatic ecosystems in NWMs are crucial for the assessment of the threat status and protection levels, the two headline indicators of the National Biodiversity Assessments (NBA), as well as the listing of threatened ecosystem types under the National Environmental Management: Biodiversity Act (NEMBA), *Act 10 of 2004* (RSA, 2004). Owing to the poor representation of inland wetlands in previous NWMs (Mbona et al., 2015; Schael et al., 2015; Van Deventer et al., 2016; Melly et al., 2016), as well as the need to improve the representation of South Africa’s estuaries, a significant effort was made in preparation for the NBA 2018, to improve the National Wetland Map version 5 (NWM5).

In addition to national obligations in the improvement of the NWM, South Africa also has international obligations in reporting the extent, biodiversity and integrity of its wetlands (inland wetlands and estuaries). The results of the NBAs are used by the Department of Environmental Affairs (DEA) to inform the global Convention on Biological Diversity (CBD), whereas the extent and quality of wetlands types are important for monitoring and reporting on the Sustainable Development Goals (SDGs) to the United Nations Environment Programme (UNEP) (UN, 2015). The extent of vegetated and inundated wetlands are, for example, required under indicators related to SDG6 (‘Ensure availability and sustainable management of water and sanitation for all’) whereas ecosystem diversity, protection and restoration is reported under SDG15 (‘Protect, restore and promote sustainable use of terrestrial ecosystems, sustainably manage forests, combat desertification, and halt and reverse land degradation and halt biodiversity loss’). The improvements to NWM5 therefore had to include both ecosystem diversity information, ecological condition, as well as cover type, including types were is inundated (lacustrine), vegetated (palustrine) or arid.

The National Freshwater Ecosystems Priority Areas (NFEPA) atlas compiled in 2011 (Nel et al., 2011) used a number of sub-national datasets to improve the spatial extent of inland wetlands mapped in NWM3, and modelled some of the wetland ecosystem types to Level 4A of the *Classification System* (Ollis et al., 2013). The NFEPA wetlands dataset had been subsequently used in the NBA 2011 for the assessment of the headline indicators of inland wetlands (Driver et al., 2012) and adopted by SANBI as NWM4. Several studies indicated that NWM4 showed up to 46 % omission errors compared to wetlands mapped at finer scales and that the hydrogeomorphic (HGM) units were incongruent between the modelled and fine-scale mapped wetlands (Mbona et al., 2015; Schael et al., 2015; Van Deventer et al., 2016; Melly et al., 2016). The shortcomings of the NWM4 could be attributed, on the one hand, to the modelling of the extent of the wetlands using space-borne *Satellite Pour l’Observation de la Terre-5* (SPOT) and Landsat multispectral imagery with inappropriate spatial resolution for detecting some of the small-scale inland wetlands in the semi-arid to arid South Africa (Thompson et al., 2002). An evaluation of the NWM4 for Gauteng, for example, revealed a commission error (where terrestrial ecosystems had been typed as inland wetlands) of 32% (Van Deventer et al., 2018a). On the other hand, the spectral bands of these multispectral imagery were unable to distinguish vegetated wetlands from adjacent terrestrial vegetation, resulting in poor representation of these systems in NWM4. Extrapolating these statistics to the rest of the country, suggested that about half of the South African inland wetlands might not be represented in the NWMs, of which two thirds are truly wetlands, resulting in only a third of South Africa’s wetlands represented in NWM4. Thus, the improved representation and typing of inland wetlands was crucial for improved assessment of the Ecosystem Threat Status (ETS) and Ecosystem Protection Levels (EPL) for the NBA 2018.

Since the NFEPA wetland maps of 2011, a number of national and sub-national wetland datasets have been published, which have enabled the improvement of the extent and wetland types across the country (Van Deventer et al., 2018a; Van Deventer et al., 2018b). The NFEPA wetlands had a tremendous impact on several levels (Nel et al., 2016). In addition, several funding sources, made available during the onset of the NBA 2018, facilitated the improvement of the extent of the wetland map in a number of district municipalities. Other examples are the National Land Cover data (GTI, 2015; 2016), which mapped the open water bodies of natural and artificial wetlands, and the Leaf Area Index (LAI), as a proxy for vegetation biomass, was generated for South Africa through inversion modelling from the Moderate Resolution Imaging Spectroradiometer (MODIS) imagery (Cho et al., 2017). In addition to the vegetation bioregions included in the National Vegetation Map update for the NBA 2018 (Dayaram et al., in prep), cover types for wetlands could be estimated. Thus several opportunities were aligned to improve the extent of the wetlands in the NWM5, the typing of inland wetland ecosystems and the determination of the cover type.

Estuaries, on the other hand, require a more accurate delineation of the spatial extent as the estuarine functional zone represents a ‘development setback line’ and feeds into supporting estuarine and coastal management processes and legislation. Since there are less than 300 functional estuaries in South Africa, it was achievable to place extensive effort into addressing previous deficiencies in the updated estuary delineation of 2018. As in the case of the inland wetlands, several projects enabled improvement in assessment of coverage of estuarine functional zones (EFZs) and the demarcation of micro-estuaries for the first time in the NBA process. An estuary is a partially enclosed permanent water body, either continuously or periodically open to the sea on decadal time scales, extending as far as the upper limit of tidal action, salinity penetration or back-flooding under closed mouth conditions (reference required). Extremes to this generic definition are during floods, when an estuary can become a river mouth with no seawater entering the formerly estuarine area or when there is little or no fluvial input, an estuary can be isolated from the sea by a sandbar and become fresh or hypersaline (modified Van Niekerk and Turpie, 2012; Van Niekerk et al., 2013). Veldkornet et al. (2015) highlighted that critical estuarine habitats, such as salt marshes and swamp forest were excluded from the NBA 2011, thereby under representing the functional zones of these systems. Additional supporting information, available since the conclusion of the previous NBA 2018 is the significant effort that has gone into updating the National Estuarine Botanical Database with field observations and more detailed mapping (Adams et al., 2016), the Light Detection and Radar (LiDAR) data that have been collected for parts of the coast, promising a spatial accuracy in mapping between 5 and 10 cm in the x, y and z spatial components, the use of supporting datasets, such as the 5 and 10 m interval above mean sea level contours (DRDLR:NGI, 2017) and the Shuttle Radar Topography Mission (STRM) 30 m data which is readily available in digital format (USGS, 2004). Estuaries are dynamic ecosystems and without a proper understanding of changes over time, these ecosystems cannot be assessed and managed appropriately. Google Earth ™ provided such a time-series dataset, allowing the mapping of the ever-changing processes that characterise SA estuaries, e.g. changes in mouth configuration and inundations patterns.

Micro-estuaries (i.e. estuaries < 500m in length and/or < 2 ha in size), river outlets, coastal seeps, ephemeral systems and waterfalls, were only represented as x,y point data in 2011, but in this iteration of the NWM an effort was also made to map some of these smaller features.

Freshwater—estuarine transition zones between rivers or inland wetlands and estuaries are defined as those areas where during any time of their hydroperiod, the system would host estuarine and riverine biota but only have abiotic characteristics of inland wetland (freshwater) ecosystems. This is defined as the freshwater—estuarine transition zone that is immediately upstream of the estuary but does not include the River-Estuarine-Interface (REI). The latter (REI) is defined as the area in an estuary within which salinity ranges from 10 to zero under the influence of the upstream limits of back-flooding or tidal intrusion. Freshwater—estuarine transition zones have previously been poorly defined, owing to their dynamic nature over long hydrological cycles and research being largely deficient with respect to their nature and characteristics (see also Rundle et al., 1998). These transition zones are essential supporting habitats to estuarine systems, and require proper mapping for the purpose of policy formulation and the protection of these zones. In addition, coastal aquatic ecosystems are also poorly understood and mapped for South Africa. A number of coastal depressions were identified according to the criterion ‘…organisms of estuarine origin (algae, crustaceans and fish, which are relicts in the case of lakes cut off from the sea since the last Ice Age) but are normally uninfluenced by the sea’ (Noble and Hemens, 1978:37).

SANBI in collaboration with the Council for Scientific and Industrial Research (CSIR) coordinated the improvement of the NWM5, supported by the participation of a number of other institutions. The aim was to improve the representation of the spatial extent and type of inland wetland and estuarine ecosystem types of South Africa in NWM5. We report the total spatial extent of inland wetlands and estuaries mapped in NWM5 as follows:

i. the extent of wetlands mapped in NWM5 in comparison to previous versions of the NWMs;
ii. the improvement in the representation of inland wetlands mapped in focus areas in NWM5 relative to NWM4;
iii. the type of cover associated with the wetlands (inundated, vegetated or arid);
iv. ecotones between rivers or inland wetlands and estuaries mapped as freshwater—estuarine transition zones; and
v. The level of confidence for the inland wetlands in terms of how well the extent and hydrogeomorphic units were captured for each sub-quaternary catchment of South Africa.

Our intention is to inform users of the improvements and shortcomings of NWM5 so that it is appropriately used in planning and decision making, whilst enabling better planning for wetland inventorying in South Africa.

## Methods

### Improving representation of inland wetlands

For inland wetlands, features mapped by the former Department of Land Affairs: Chief Directorate of Surveys and Mapping (DLA:CDSM, 2006) which had been incorporated in NWM4 were extracted and retained for use in the NWM5. These included all types of pans, river areas, lakes, marshes and vleis. This dataset was readily available at a national scale, since it was merged and used in mapping of the NFEPA wetlands (NWM4). The other updates from the now Department of Rural Development and Land Reform: Directorate National Geo-Information (DRDLR:NGI) of 2009 and 2012 were, however, not yet merged and topologically cleaned for use at a national scale. An updated version of these hydrological features, were collected from the DRDLR:NGI (2016) as provincial geodatabases at the end of March 2016. Hydrological features related to inland wetlands included dry, salt, non-perennial and perennial pans, water course features, flood banks, lakes, marshes or vleis, mudflats, pools, river areas and swamps. The DRDLR:NGI MapInfo provincial geodatabases were imported and merged into a single feature class in ArcGIS 10.3 (ESRI, 1999-2014). The data were projected to the South African coordinate system used by the NBA 2018, the Albers Equal Area (AEA) Conical projection with the spheroid and datum being the World Geodetic System of 1984 (WGS84). This coordinate system least distorts the surface area extent calculated for ecosystems. This coordinate system uses the 25°E as central meridian with two standard parallels including 24°S and 33°S. The topology was cleaned to avoid duplicate or overlapping polygons, and subtypes were defined to enable consistent distribution mapping by multiple data capturers across the country.

Firstly, all nationally available datasets were incorporated, including the Working for Wetlands data available from the Biodiversity Geographical Information System which were mapped by SANBI since 2006, peatlands data from the Water Research Commission (WRC) Report No. 2346/1/17 (Grundling et al., 2017) and the extent of the estuaries mapped for the NBA 2011 (Van Niekerk and Turpie, 2011; Van Niekerk et al., 2013). The spring points from the DRDLR:NGI 2016 dataset were buffered by 2 m, and classified as seeps. The pans and river areas, mapped in 2006 and 2016 by DLA:CDSM and DRDLR:NGI respectively, were translated to Level 4A (HGM units) of the *Classification Systems* as depressions and rivers. HGM units assigned by the Working for Wetland teams were kept as is. All feature names were corrected and the version was called the NWM version 5.2. NWM5.2 was clipped to provinces and distributed to data capturers.

Secondly, all available fine-scale datasets (see Van Deventer et al., 2018a; Van Deventer et al., 2018b) were merged with the clipped version of NWM5.2, into version 5.3. A merge was used to easily identify overlapping areas where the data capturer then evaluated the multiple, overlapping polygons from diverse studies and judged which one should be retained, if not all. Available inland wetland data were integrated and additional wetlands were mapped by data capturers for nine focus areas (Table 1) for the period between 1 September 2016 and 31 March 2017. Datasets from three other study areas also improved the NWM5 draft versions. These were the West Rand District Municipality (USAID, 2018) and two study areas from the WRC K5/2545 project, the southern part of the sub-quaternary catchment (SQ4) 7439 (in quaternary catchment S32D) around the town Hogsback and the western part of the SQ4 1375 (in quaternary catchment W55A) in which Tevredenpan is situated (Van Deventer et al., 2017).

**Table 1:**
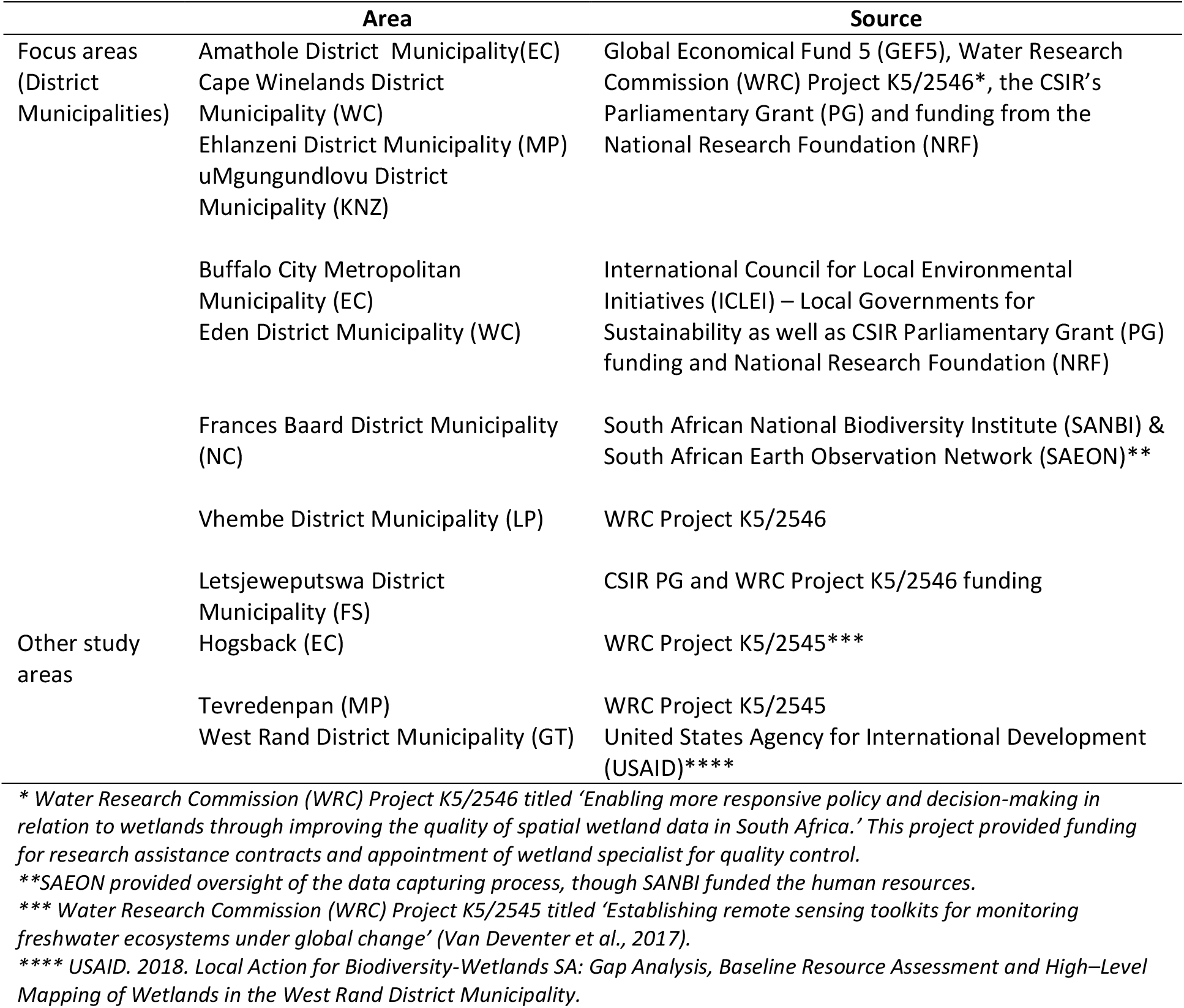
Focus areas and other study areas where mapping of HGM wetland types were improved and included in NWM5, and associated funding sources.

During the integration and mapping phase, the attributes of inland wetland types had to be completed for Levels 1, 3 and 4A of the *Classification System*. Back-drop imagery included the freely available 50 cm colour orthophotography available through the ArcGIS online web map service from DRDLR:NGI dated from 2012 to 2013. SPOT imagery was also used in some instances, dated to similar years. Unfortunately, most of these images were largely taken during the dry season, possibly to avoid cloud cover, and were therefore less suitable for the purpose of mapping of wetlands. Where the data capturer found it difficult to judge the extent or HGM type of the wetland, historical images available through Google Earth ™ were accessed to support the mapping of inland wetlands.

Decisions regarding the extent and ecosystem type for features were guided primarily by three principles: to always map the maximum extent of a wetland, if possible to map the original extent (historic maximum), and to retain the extent and typing done in fine-scale datasets. The focus areas were then reviewed by national wetland experts (Freshwater Consultancy Group Pty Ltd (FCG) and Wetland Consultancy Services Pty Ltd (WCS)) and corrections implemented by data capturers. Subsequently the available data for the remainder of the provinces were integrated with limited mapping of large floodplains, Ramsar sites and nine limnetic depressions. Limnetic depressions were considered unique wetland types where the maximum depth exceeds 2 m at the average annual low-water level of an open waterbody (Ollis et al., 2013). The nine limnetic depressions mapped in NWM5 included Barberspan (North West), De Hoop (Western Cape), Groenvlei (Western Cape), Lake Banagher, Lake Chrissie and Tevredenpan (Mpumalanga), Lake Fundudzi (Limpopo), Lake kuZilonde (KwaZulu-Natal), and Lake Sibaya (KwaZulu-Natal) (compiled from Hill, 1969; Miller, 1998; Noble and Hemens, 1978; https://www.lakepedia.com/). Following the integration of the provincial datasets, these were reviewed and edits implemented.

### Mapping artificial wetlands as a separate layer

To better represent the original wetland extent for the NBA 2018, artificial wetlands were compiled as a separate feature class layer in the ArcGIS geodatabase. This was to assessdisturbance on wetland ecosystem types in the NBA 2018 assessment report. Artificial wetlands were compiled from the DLA:CDSM 2006 dataset included in NWM4, and from updated hydrological data received from DRDLR:NGI in 2016, the large dams register (approximately 159 dams) dataset from the Department of Water and Sanitation (DWS, 2015), as well as farm dams mapped for the DWS Verification and Validation project for selected tertiary catchments in the Breede- Gouritz Water Management Area and the KwaZulu-Natal Province (DWS, unpublished results). Overall, the features included in this dataset are large, closed and open reservoirs, large state dams, smaller farm dams, fish farms, pools, purification plants, sewerage works, slimes dams, tailing impoundments and water tanks. Where artificial wetlands were situated within an inland wetland ecosystem type, the artificial wetland polygon had to be merged with the adjacent polygon to represent the original extent prior to the modification. Isolated artificial wetlands were completely deleted from the NWM5.

### Improving representation of estuaries

The mapping convention for determining the Estuary Functional Zone (EFZ) was based on a precautionary approach with the departure point being the first inland 5 m contour above mean sea level, to capture all estuarine processes and biotic responses. This dataset was then adjusted to address the shortcomings identified by Veldkornet et al. (2015), e.g. exclusions of swamp forest (fresh water mangrove) and salt marsh areas contiguous to estuaries. In addition, all habitat features excluded from ground-truthed vegetation maps were also incorporated in the new delineation (Adams et al., 2016). The EFZ boundary, where possible, mapped the maximum extent of the historical and present geographical boundary (although resources were only available to do this accurately in KZN). The location of estuary outlets was determined by the maximum extent of migration of the estuary mouth or outlet (furthest north and south) as identified from any historical image, i.e. Google Earth ™ or historic aerial photographs. Areas identified by coastal LiDAR datasets (corrected to mean sea level and verified by experts) or the 1:100 year flood line delineation (only available for the Groot Berg or Breede estuaries) were also incorporated. All ‘island’ type features created by high elevation areas surrounded by estuarine floodplains were incorporated into the EFZ. In addition, in the case of small, incised estuaries (i.e. where the 0 m, 5 m and 10 m contour are close to one another) with relative high river inflow, delineation was based on the 10 m contour above mean sea level to accommodate mapping uncertainty and lateral movement. The EFZ was also extended to incorporate environment that is predominantly surrounded by estuarine habitats or processes (i.e. more than 75% of feature is surrounded such as s-bends or oxbow bend). In addition, habitat features that support estuarine functioning were also included to ensure future health, i.e. upstream inland wetlands that influence estuarine water quality by filtering nutrients. This included the incorporation of small areas of inland aquatic ecosystems contiguous to estuaries e.g. seeps and springs. Overall, the 2018 revised EFZs strived to incorporate all vegetation ecotones that have elements of estuarine habitat, e.g. mosaics of swamp and dune forest. The EFZ was broadened to include novel ecosystems such as marinas and harbours adjacent to estuaries as they directly influence estuarine functionality and biodiversity. Where possible, names were changed to provincial standards, e.g. the KwaZulu-Natal (KZN) provincial gazettes and Eastern Cape conservation plan.

Estuarine and provincial inland aquatic datasets, now including inland wetlands and the extent of some large rivers, were merged into a national dataset. The draft NWM5 was checked for overlapping polygon (topology) errors, features aligned along provincial boundaries as well as between inland aquatic ecosystems and estuaries. The total extent (hectares) of all the NWM versions 1 to 5 were updated in ArcGIS and calculated as a percentage of the total of South Africa’s extent. Similarly, the surface area of each estuarine and inland aquatic ecosystem type is summarised and the percentage calculated for the total extent of South Africa. These statistics of NWM4 and NWM5 are then reported for the ten focus areas to highlight the improvements of the digitising efforts.

### Determining inundated, arid and palustrine wetland extent for SDG reporting

To determine the extent of inundated wetlands, the seasonal and permanent water and wetland classes from the 30 m National Land Cover data of 2013/4 (GTI, 2015) were extracted. The extent of the inundation was then calculated as a percentage of the extent of all inland wetlands. Subsequently, the remaining extent of the inland wetlands were combined with the LAI predicted using the 463 m spatial resolution MODIS image of 13 March 2013 (Julian day 73) to indicate potential ranges of vegetation biomass (Cho et al., 2017). A LAI range of 0 – 1 was used to distinguish low to no vegetation and therefore likely to be arid, whereas a LAI range from 1 to 8 was considered to indicate dense grass to tree cover, and therefore more likely to be palustrine wetlands (Prof Moses Cho, *pers. comm*.). The total amount of pixels for both processes was extracted using the zonal statistics tool in ArcGIS 10.3 (ESRI, 1999-2014).

### Mapping freshwater—estuarine transition zones

In addition to the EFZs, the freshwater—estuarine transition zones between the rivers or inland wetlands and estuaries were also defined, namely as river reaches where only riverine abiotic processes dominate, but also where riverine and estuarine biota occur (during any time of their hydroperiod). These areas are not subjected to tidal action or back flooding and at no stage experience an increase in salinity as a result of tidal penetration. These transitions are poorly understood and have previously not been delineated, owing to their dynamic nature over long hydrological cycles. River lengths, varying between 0.5 km and 30 km, were defined based on sampling data, topography and expert opinion. The steeper the gradient, the smaller the transition zone. Thus rivers with extensive lowland reaches such as the Breede have longer transition zones. Ephemeral rivers upstream of estuaries were excluded from this delineation. The extent of the freshwater—estuarine transition zone were mapped and included in the national rivers database of the NBA 2018 (Smith-Adao et al., 2018).

Freshwater—estuarine transition zones have been identified based on expert knowledge of the occurrences of estuarine-associated fish and invertebrate assemblages. These assemblages within the freshwater—estuarine transition zones are a mixture of typically estuarine species e.g. moony *Monodactylus falciformis*, those equally adept at completing their entire lifecycle in both habitats e.g. estuarine roundherring *Gilchristella aestuaria*, freshwater species that may have an estuarine phase of their life-history e.g. multi-specific freshwater prawns of the Genus *Macrobrachium* and catadromous eels Anguillidae that are either resident, passing through whilst recruiting to the upper catchment or migrating back as adults to spawn far offshore in the abyssal depths or freshwater swimming crabs *Varuna litterata* that are spawned and require early development at sea before recruitment back to the freshwater—estuarine transition zones. Typically estuarine species include, but are not limited to, ‘facultative catadromous’ fish such as freshwater mullet *Pseudomyxus capensis* that may opportunistically spend most of their life in freshwater but return to spawn in estuaries or the sea. Appendix II lists the extent of river transition zones, where these species assemblages have been observed.

Freshwater—estuarine transition zones were not incorporated in to the EFZ, as they are not subject to estuarine abiotic processes, but should be highlighted as estuarine supporting areas in current planning legislation and approaches to ensure that future developments/ discharges/ abstraction/ infrastructure do not disrupt or degrade estuarine connectivity and ultimately condition. Coastal depressions were refined from Noble and Hemens (1978) using available grey and scientific literature, as well as expert opinion (Van Deventer et al., 2018). The limnicity status of these systems has been added to the NWM5 database under fields related to the hydroperiod. Other inland wetlands located on sandy coastal plains near the coast or estuarine systems, were also attributed as ‘coastal’ in the NWM5 dataset.

### Confidence ranks of the inland wetlands

The final step was a confidence map, generated for the inland wetlands based on the extent of a sub-quaternary catchment (SQ4) that was mapped in full, as well as the degree of expertise involved in the wetland mapping, the completeness of the HGM unit and extent to which the hydroperiod is known for a wetland (Table 2). Statistics are reported for the number of SQ4s relative to the total number of SQ4s in South Africa for each of the five ranks, as well as the percentage of surface area of South Africa which is likely to be mapped according to the ranks.

**Table 2:** Confidence ratings assigned to sub-quaternary catchments based for inland wetlands.

| Rating | Description |
| --- | --- |
| 1 – Low | Desktop mapping of the extent of inland wetlands was done by non-wetland specialists for a part of, or to the full extent of the SQ4.<br>(Mostly DRDLR:NGI for non-wetland typing purposes) |
| 2 – Low to Medium | Desktop mapping of inland wetland extent was captured by interns for the purpose of NWM5 to the full extent or a part of the SQ4. This may also include areas where data from wetland specialists had been incorporated, but the dataset was either not typed to the HGM unit, or complete for all HGM units, or was based on old imagery. |
| 3 – Medium | Desktop mapping of the extent of inland wetlands and HGM typing was done by wetland specialists for the full extent of the SQ4. |
| 4 – Medium to High | Desktop mapping of the extent of inland wetlands and HGM typing as well as field verification and revision by experts was completed for the full extent of the SQ4. |
| 5 – High | Inland wetlands have been mapped and verified for a period of > 10 years over multiple hydrological cycles / hydroperiod for the full extent of the SQ4.<br>Verification may include field observations as well as soil and/or vegetation surveys. |

## Results

### Improvement of the National Wetland Map 5 compared to previous versions

A total of 4 698 824 ha of inland aquatic ecosystems and artificial wetlands have been mapped in South Africa, constituting about 3.9 % of the surface area of the country^1^ (Table 3; Figure 1; Appendix I2). Aquatic ecosystems, including estuaries, inland wetlands and some river channels, totalled 4 100 434 ha (3.4%), while wetlands (inland wetlands and estuaries) totalled 2 824 767 ha (2.3%). The extent of the ecosystems and attributes represented in NWM5 has increased compared with the previous versions of the NWMs (Figure 3). In addition, the artificial layer has been separated from the inland aquatic data, and now forms part of a collection of datasets in a geodatabase called the South African Inventory of Inland Aquatic Ecosystems (SAIIAE) (Van Deventer et al., 2018a; Van Deventer et al., 2018b).

**Table 3:**
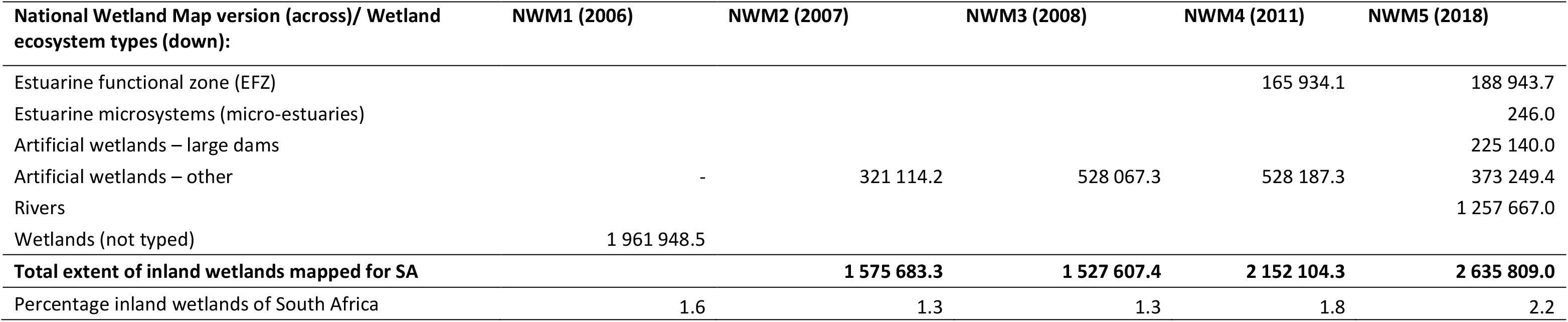
The spatial extent of natural and artificial aquatic ecosystems represented in the National Wetland Map versions 1- 5. The full extent of the NWM5 is reported in Van Deventer et al., 2018a, whereas the full extent of estuaries, including their off-shore extent, is reported in Van Niekerk et al. (2018)

**Figure 1:**
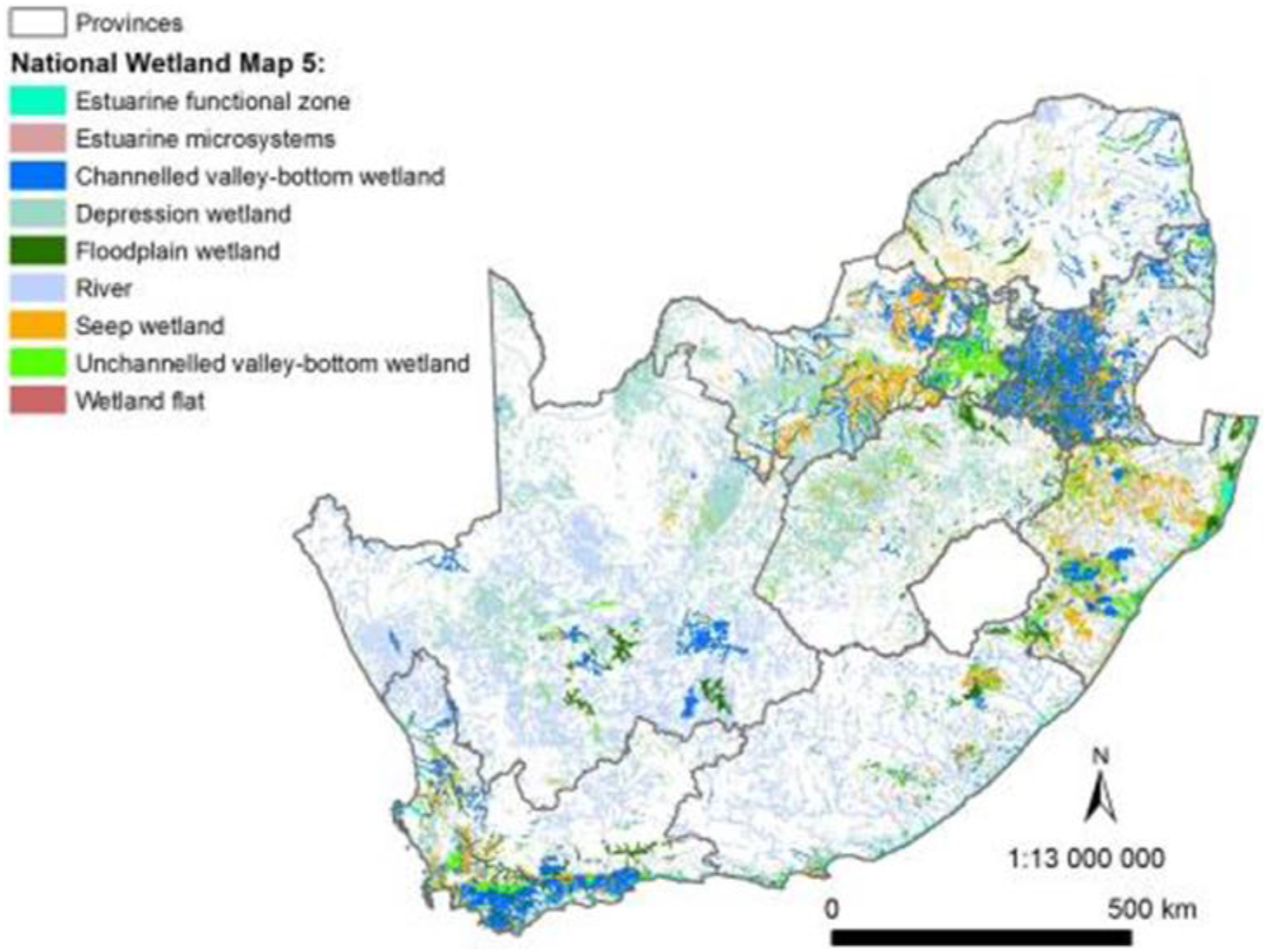
The improved representation of inland wetland and estuarine ecosystems of South Africa in National Wetland Map 5.

In the NWM5, inland wetlands constituted more than 147 000 polygons, totalling more than 2,6 million ha or 2.2% of South Africa (Table 3). The inland wetlands have increased in extent from NWM4 (NFEPA wetlands of 2011) to NWM5 by 123%; and now comprising 2.2% of the surface area of South Africa (Figure 2). The extent of the 292 EFZs in NWM5, which falls within the boundary of South Africa increased by 121%, to 0.15% from the previous version where 0.14% of the country’s surface area constituted estuaries (Table 3), although this may increase further as the off-shore (marine transition) is reported (Van Niekerk et al., in prep). Forty-two microsystems (micro-estuaries) have been added in NWM5, which represents 213 ha of systems along the coast. The representation of artificial wetlands increased by 113% from 528 187 ha in NWM4 to 598 389 ha in NWM5, making up 0.5% of South Africa’s surface area. Artificial wetlands show a minor overlap with natural inland systems of 37 172 ha or 0.03% of SA. Large dams comprise 0.18% of the surface area of the country, and the remaining artificial wetlands 0.3%.

**Figure 2:**
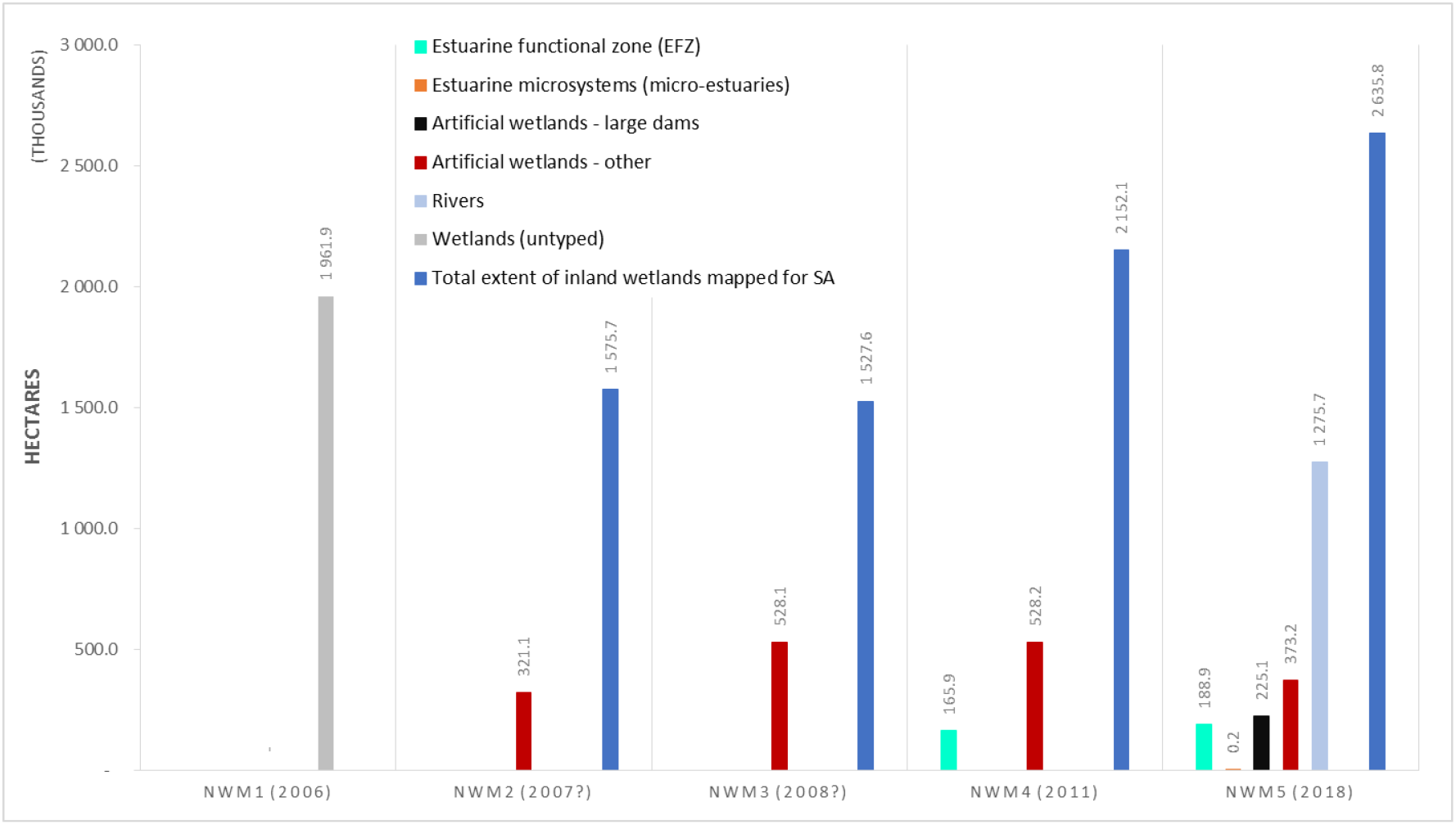
The extent of wetland ecosystem types represented in the first five versions of the National Wetland Map of South Africa.

### Improvement in the spatial extent of inland wetland ecosystems per HGM type

Depressions were the HGM unit with the highest percentage of representation relative to the surface area of South Africa (764 739 ha; 29% of the extent of inland wetlands), followed by Channelled valley-bottom systems (671 346 ha), Floodplains (542 819 ha), Seeps (453 748 ha) and Unchannelled valley-bottom wetlands (187 891 ha) (Table 4). Wetland flats showed the lowest percentage of representation (15 267 ha), comprising only 0.6% of the spatial extent of inland wetlands mapped in NWM5, and 0.01% of the extent of the country’s surface area. The majority of these are located in the Western Cape Province (80%, results not shown here), with 12% of the wetland flats mapped in the Northern Cape Province, 6% in the north-western parts of the Free State Province, and < 2% mapped in the Eastern Cape, KwaZulu-Natal and Limpopo Provinces combined.

**Table 4:**
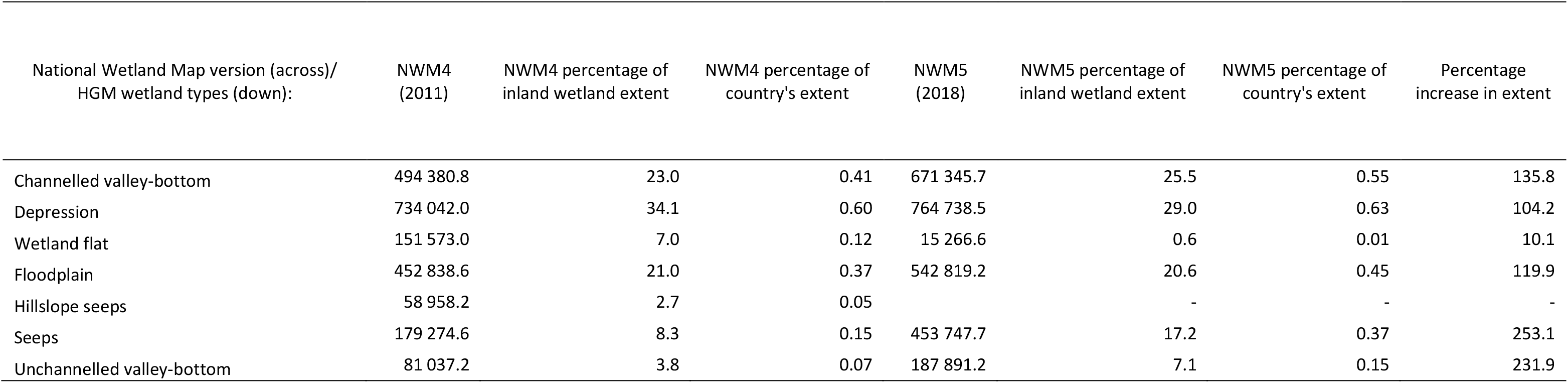
Spatial extent and percentage of the hydrogeomorphic (HGM) units relative to the total extent of inland wetlands and the surface area of South Africa.

When comparing the spatial extent of the HGM units of NWM5 with NWM4, all HGM types showed a marked increase in extent of > 120%, except the wetland flats (Table 4; Figure 3). Wetland flats modelled in NWM4 were corrected to Depressions in NWM5 resulting in a marked decrease of 90%. Hillslope seeps have been amalgamated into the Seeps category, resulting in a 253% increase in the spatial extent, when comparing the combination of Hillslope seeps and Seeps of NWM4, to Seeps of NWM5 (Figure 3). When the percentage of HGM units are compared between NWM4 and NWM5, relative to the total extent of inland wetlands mapped, NWM5 mapped more Valley-bottom and Seep systems (3-9% more in the extent) compared to NWM4. The extent of Depressions, Wetland Flats and Floodplains were greater in NWM4 compared to NWM5 (0 – 6% more in extent). It is interesting to note that Floodplain wetlands achieved a similar percentage (21%) of the total spatial extent of inland wetlands.

**Figure 3:**
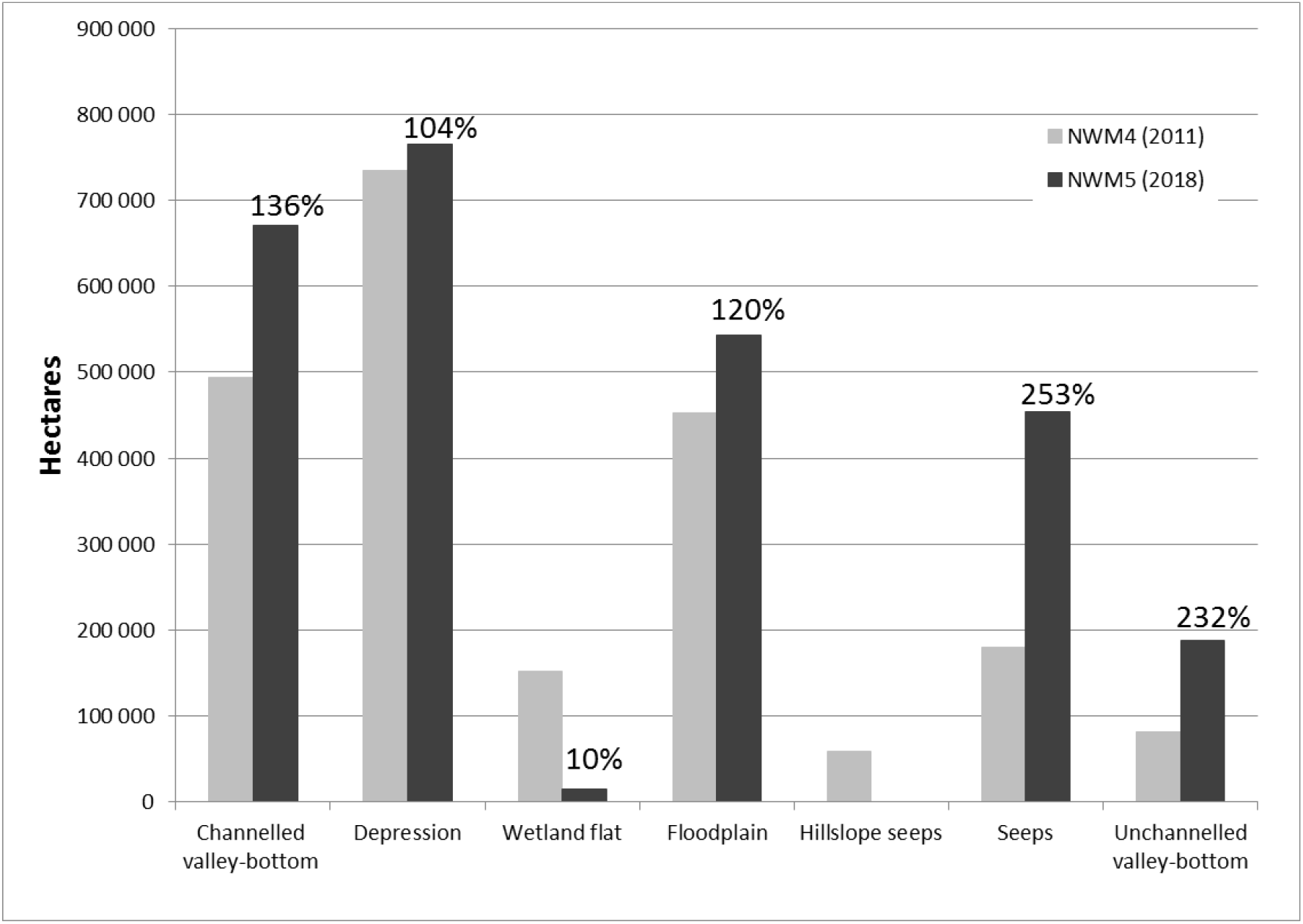
Comparison of the spatial extent of hydrogeomorphic (HGM) units between NWM4 and NWM5. The percentage change (5) is indicated above the relevant bar of NWM5.

### Improvement of the representation of inland wetlands in focus areas

The majority of the ten focus areas, where wetlands data were integrated and additional wetlands mapped in NWM5 for the NBA 2018, showed an increase in the extent of inland wetlands compared to NWM4 (Table 5; Figure 4). Five of the ten focus areas (Amathole, Buffalo City, Cape Winelands, Lejweleputshwa and the Frances Baard municipalities), however, showed a reduction in the extent of inland wetlands. For these areas, commission errors from remote sensing and probability mapping, which had been included in previous versions of the NWMs, were removed in NWM5. The four focus areas (Eden, Ehlanzeni, Vhembe and the West Rand District municipalities) showed increases in the extent of inland wetlands of between 154% and 227%. The West Rand District Municipality achieved the largest increase in extent of nearly 700% compared to NWM4. The extent of inland wetlands for the Hogsback and Tevredenpan study areas increased from the NFEPA (NWM4) wetlands to NWM5 by 207% and 390% respectively, following in-field visits and corrections (Table 5). The total extent of inland wetlands ranged from <1% to 43% of the surface area of the respective study areas.

**Table 5:**
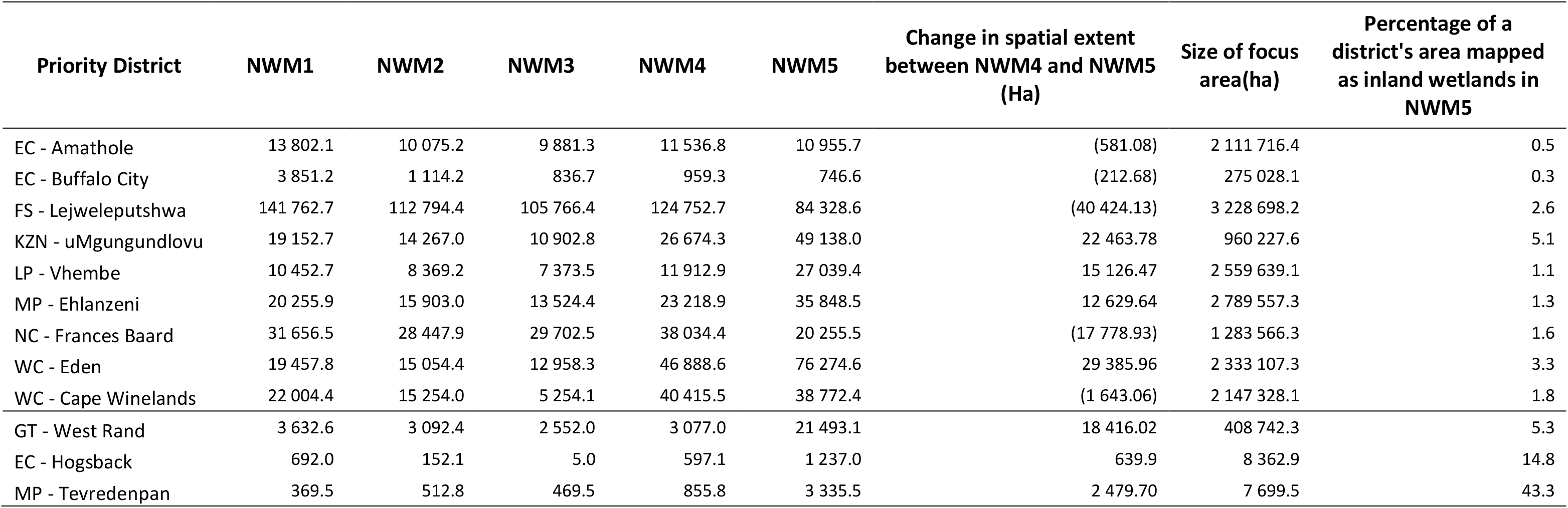
Extent of inland wetlands (in hectares) mapped in NWM5 and former NWMs for focus areas as well as the percentage (%) it constitutes of the surface area of the district or study area.

**Figure 4:**
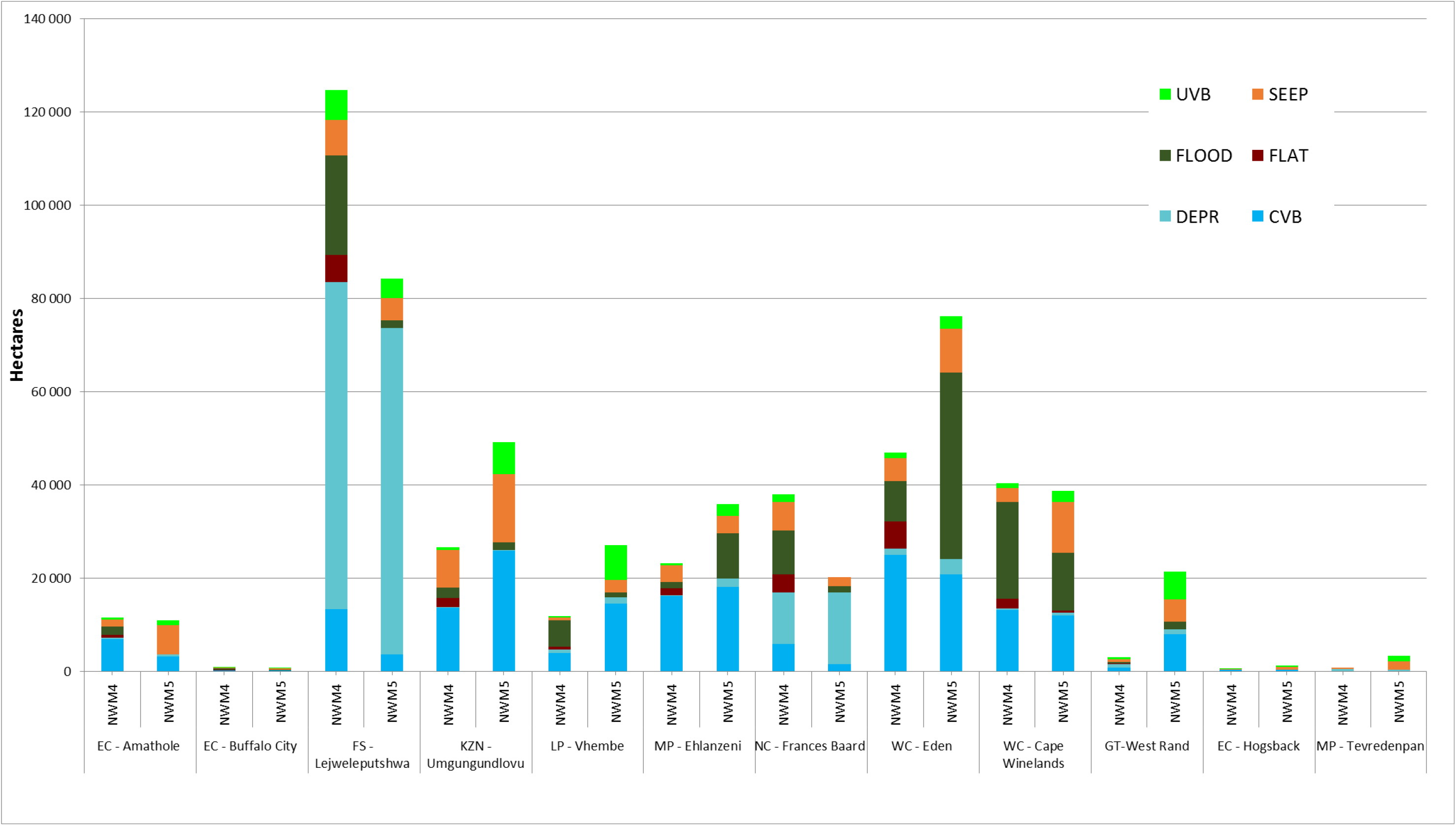
Comparison between wetland ecosystem types for focus areas. *Acronyms used: EC = Eastern Cape; FS = Free State; GT = Gauteng; KZN = KwaZulu-Natal; LP = Limpopo; MP = Mpumalanga; NC = Northern Cape; NW = North West; WC = Western Cape.*

### Cover types of inland wetlands (inundated, palustrine or arid) for SDG reporting

Inundated wetlands made up an estimated 278 719 ha or 11% of the extent of South African inland wetlands (Figure 5) and are distributed across the country. Inland wetlands which are more likely to be vegetated or palustrine (LAI > 1) are found in the Fynbos, Grassland and Savanna Biomes of South Africa, totalling an estimated 1 447 932 ha or 55%. Arid systems are located in the central Karoo and Northern Cape Provinces primarily, and a total of 909 157 ha or 34% have been estimated where the LAI < 1.

**Figure 5:**
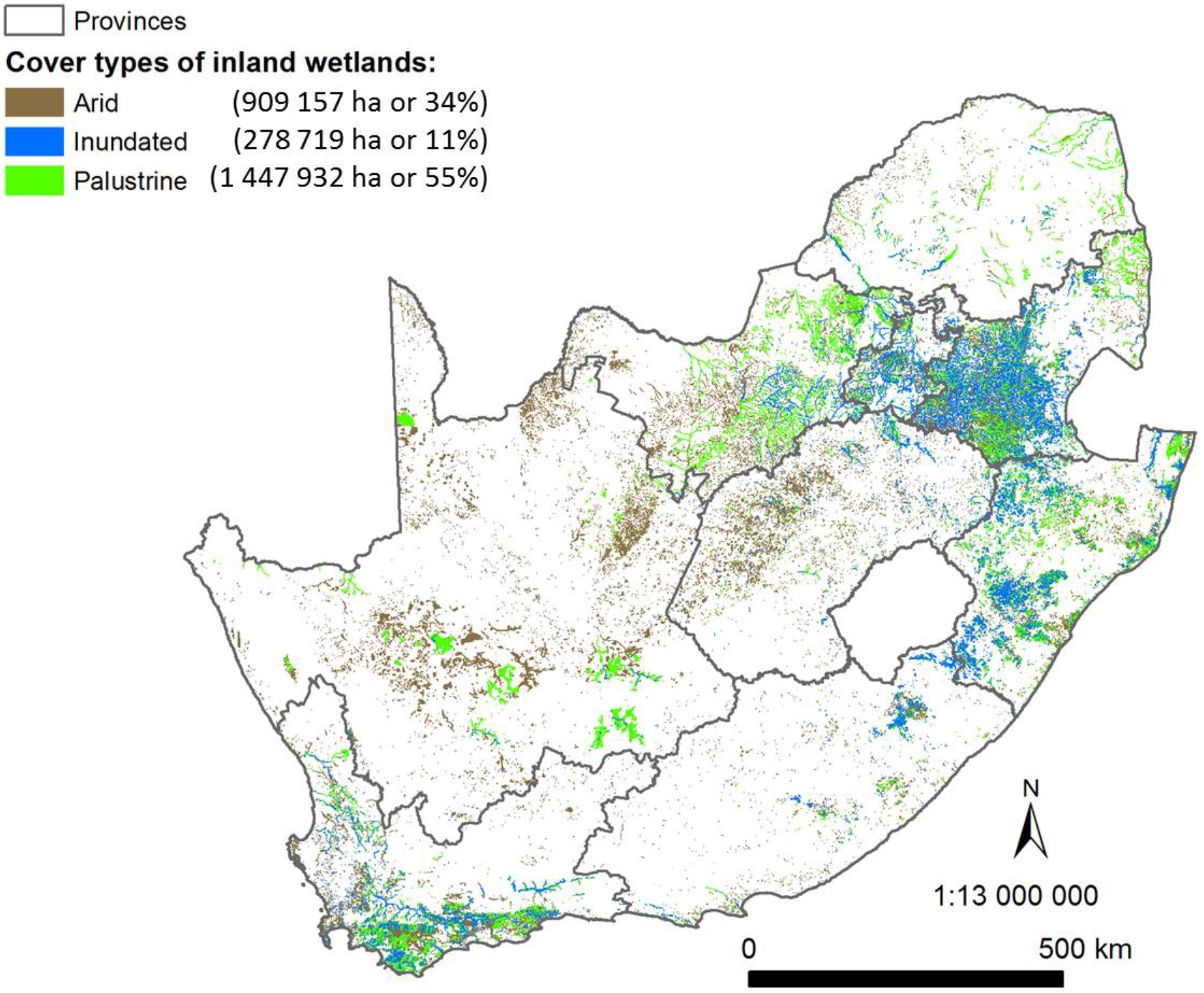
Distribution of cover type of inland wetlands across South Africa.

### Estuarine support areas (freshwater—estuarine transition zones of rivers) and coastal depressions

A total of 1 931 km of transition rivers (Figure 6) have been identified (Appendix II) which constitutes about 1% of the total length (164 018 km) of rivers as identified by Smith-Adao et al. (2018). Almost 30 000 ha of inland wetlands were found to coincide with the coastal regions of South Africa (Table 6). The majority of these are coastal depressions, of which Groenvlei (357 ha) and Lake Sibaya (8 233 ha) were the only limnetic depressions.

**Table 6.** Extent (ha) of coastal systems as mapped in National Wetland Map 5.

| Hydrogeomorphic units | Hectares | Percentage of all inland wetlands in the coastal region of South Africa |
| --- | --- | --- |
| Channelled valley-bottom systems | 1 667.3 | 5.7 |
| Depressions | 22 771.4 | 77.7 |
| Wetland flats | 16.7 | 0.1 |
| Floodplains | 558.0 | 1.9 |
| Seeps | 2 397.9 | 8.2 |
| Unchannelled valley-bottom systems | 1 908.8 | 6.5 |

**Figure 6:**
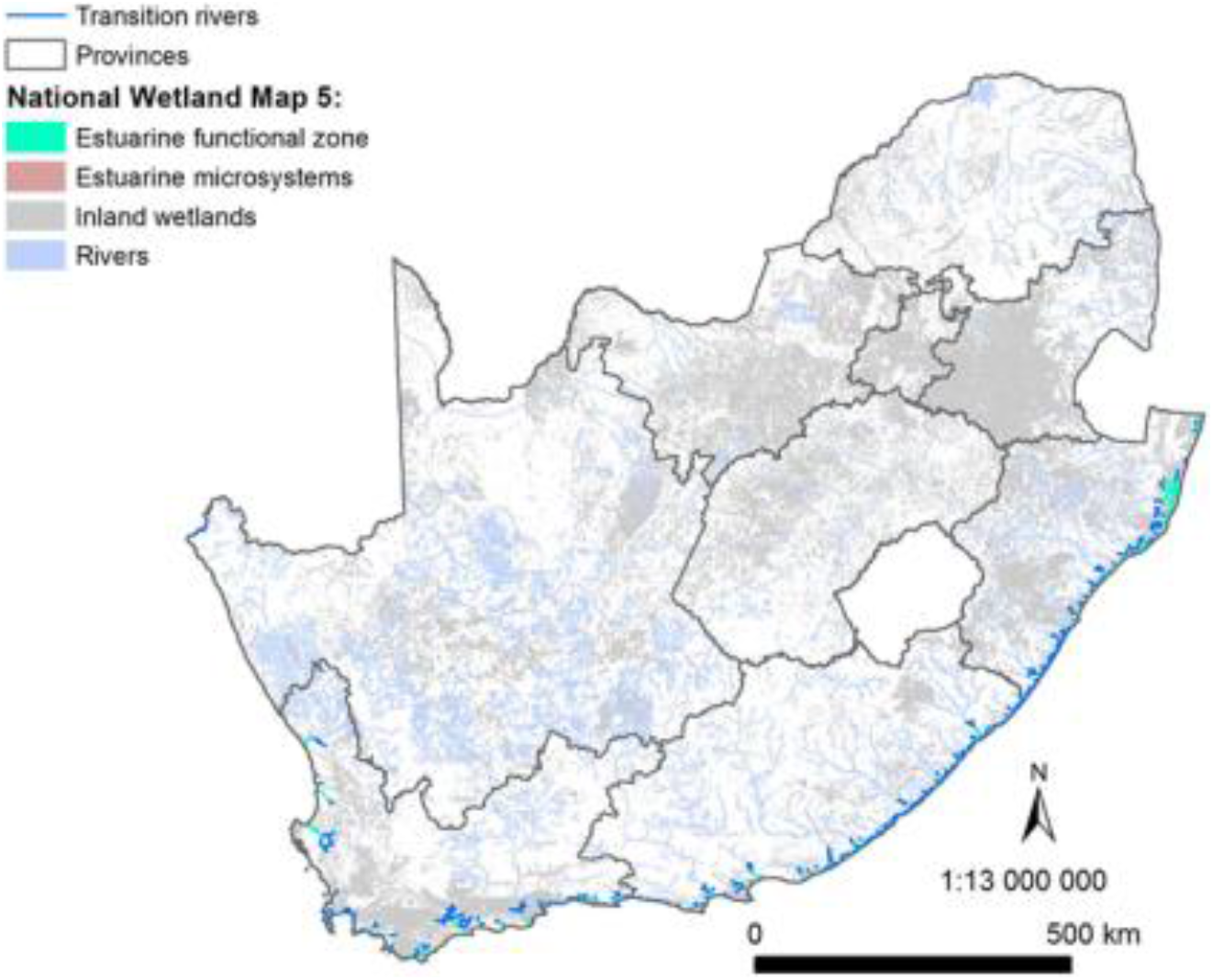
Estuarine support zones (freshwater—estuarine transition zones of rivers).

### Confidence ranking of the inland wetlands of NWM5

The majority of the country has been mapped by non-wetland experts with limited understanding of wetlands with a low confidence overall (76% of country’s surface area, Figure 7, Table 2). Almost 17% of the surface area of the country has been attended to at a desktop level through the integration of existing data and/or the mapping of wetlands by interns trained during the update of NWM5 (representing low-medium confidence). Only 7% of the country has been mapped and typed to HGM units by wetland experts (i.e. medium confidence), and a further 0.04% of the country including site visits and subsequent improvements to the representation (extent and ecosystem typing) of the map. No area has been mapped and refined following long-term research (category 5 = 0%) (i.e. high confidence).

**Figure 7:**
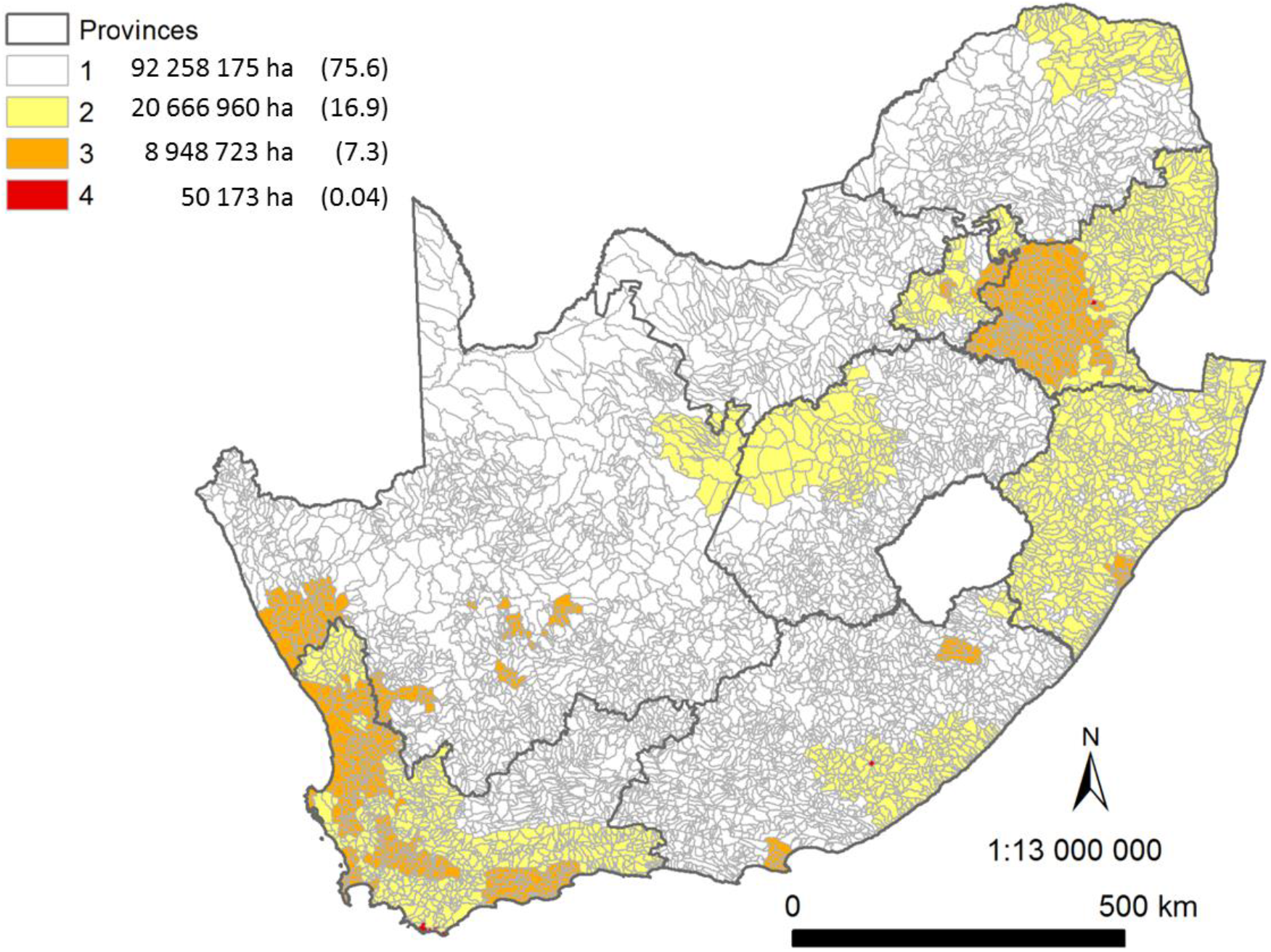
Areas of confidence in the spatial extent and HGM units for inland wetlands. Categories include: 1 – Low; 2 – Low to Medium; 3 – Medium; 4 – Medium to High; and 5 – High. The extent (ha) and percentage of the country’s surface area is indicated in brackets after each category.

## Discussion

In the past four years, a significant effort has been made to improve the representation of various datasets in the South African Inventory of Inland Aquatic Ecosystems (SAIIAE), particularly the National Wetland Map version 5 (NWM5) in preparation for the National Biodiversity Assessment 2018, and for reporting to international conventions on behalf of South Africa. The effort took more than 30 data capturers from more than 10 organisations, at a total estimated cost of R7 million to generate (Van Deventer et al., 2018). The results paid off, in that a total amount of 4 698 823.6 ha (3.9% of South Africa) of inland aquatic ecosystems and artificial wetlands have been mapped for South Africa. Yet, despite this tremendous effort, the majority of sub-quaternary catchment extents were at a low confidence that the extent and hydrogeomorphic unit have been well represented at a desktop level. Wetlands in arid to semi-arid regions are poorly detected through remote sensing indices, which often use open water indices for extracting wetland extent. As a result, few of the palustrine and arid systems are well represented in regional to global wetland maps. The extent of wetlands for Africa, for example, has been estimated at 22 440 000 ha or 0.7% of the surface extent of Africa in the Global Lakes and Wetlands Database (GLWD) (Lehner and Döll, 2004). The more recent Global Inundation Extent from Multi-Satellites (GIEMS) dataset (Fluet-Chouinard et al., 2015), at a spatial resolution of 420 × 460 m (total about 19,3 ha per pixel), estimates the extent of wetlands in South Africa at 4.2% of the surface area. It does not, however, distinguish between natural or artificial wetlands, or amongst estuaries, rivers or inland wetlands. Previous wetland maps for South Africa, done using remote sensing and prediction modelling, have proved to underrepresent the full extent of arid and palustrine wetlands. In semi-arid to arid countries, such as South Africa, heads-up digitising and in field verification is essential to improve our national maps.

The investment made in the NWM5 showed that a significant improvement in the representation of inland wetlands can be achieved. Commission errors associated with the extent of wetlands were removed for some focus areas, resulting in a decrease of the incorrect representation of inland wetlands. Several focus areas, however, showed an increase in the extent of wetlands. During the update of the NWM5, the base data of wetlands for the West Rand District Municipality was combined with the *wetlands probability map* of Collins (2018), resulting in an increase in extent by nearly 700%, though without in-field verification, the confidence ranges between low and moderate. In-field verification of wetlands by wetland ecologist (Mr Anton Linström and Ms Nancy Job) for the Tevredenpan and Hogsback study areas, resulted in an increase of more than 200% and a moderately high confidence rank (Van Deventer et al., 2017). Investment in similar efforts is therefore crucial for improving the representation of wetland extent to a moderate confidence level. Further improvements should be done for selected priority areas, at catchment levels, particularly in the strategic water source areas (Nel et al., 2017; Le Maitre et al., 2018), areas of high development pressures, and others identified through the NBA 2018 assessment of threatened ecosystems.

Estuaries improved by less than 5% from previous version, with some smaller systems moved to the micro-estuaries category and 4 new small estuaries being mapped and added. However, significant improvements were made in the incorporation of all estuarine and estuarine associated habitats in the updated delineation. In addition, the dataset now includes 42 micro-estuaries. The use of time series data vastly improved the mapping of the estuarine extent and allowed for the incorporation of dynamic features such as the river or estuary mouth position. LiDAR data showed significant promise, but unfortunately the datasets were not post-processed adequately, e.g. included the return signal from tree tops instead off ground level, to be used without expert judgment. In future, this type of dataset would benefit from extensive post-processing to increase reliability and assist with increasing mapping accuracies. The incorporation of 1:100 year flood lines allowed for the incorporation of all relevant sediment processes, but was unfortunately only limited to two large systems. However, they supported the use of the 5 m contour above mean sea level as a proxy for sedimentary and inundation processes. While the 5 and 10 m above mean sea level contour dataset (DRDLR: NGI, 2017) proved to be useful in supporting the delineation of estuaries in the lower reaches where the floodplain opens up on the coastal plain, it was less useful for delineating the incised, upper reaches correctly, e.g. the Palmiet was delineated a third shorter than the measured extent. It is clear that use of expert judgement in combination with a precautionary approach is advisable in delineating the upper reaches of most systems with limited extent. Estuary delineation in South Africa is still largely based on spatial and habitat features as detailed information on soil moisture, sediment particle size, redox potential, and total organic matter is not available on a national scale. The latter have been used in international approaches on regional scale delineations and therefore highlight future research requirements (Adam, 1992; Caeiro et al., 2003; Junk et al., 2013).

Although the index on SDG6, ‘Ensure availability and sustainable management of water and sanitation for all’, requires only the reporting of lakes (the open water) and vegetated wetlands at a national scale, we propose the inclusion of arid systems which are not permanently or seasonally inundated, nor vegetated as an additional category. The results showed that < 20% of South Africa’s inland wetlands are inundated, and of the remaining wetlands, 55% could be vegetated and 34% arid. Both the land cover (GTI, 2015) and LAI (Cho et al., 2017) products, at 30 m and 463 m spatial resolution, are unfortunately at a very coarse spatial resolution and considered inadequate to accurately determine the true extent of inundation and vegetation cover of inland wetlands. Closer inspection of sites showed an underrepresentation of the inundated waterbodies, and over-estimation of the vegetated and arid systems. Finer-scale data or in-field verification is therefore required to verify the extent of inundation and vegetation of wetlands, whereas in-field verification of the limnicity of systems is crucial. Time-series analysis should be incorporated to determine the full hydroperiod and phenology of wetlands. These results are therefore only potentially broad-scale indicators of inundation, vegetation and arid systems for the purpose of reporting at a national scale.

Demarcating the freshwater—estuarine transition zone highlights the importance of the river reaches just above estuaries and the need to consider their role in maintaining estuary condition. In the past five years there has been a trend to move/consider moving new waste water discharge points out of estuaries to just above the estuary to allow for the application of less stringent standard / special standards license agreements to river discharges versus the more onerous requirements of the receiving environment applicable to estuaries. Benefits of these transition zones include refuge from adverse conditions such as hypersalinity, eutrophication, hypoxia and temperature extremes in the estuarine environment. Similarly, pathogens and parasites that latched on in estuaries and the sea may be shed through osmotic stress in these transitional freshwater reaches. Fish and invertebrates can also benefit from a greater diversity and abundance of grazing and prey that may be unavailable or limited in the adjacent estuarine habitat. In turn, lower predation levels arise from stenohaline estuarine predators not being able to follow euryhaline prey. Freshwater—estuarine transitional zones also provide extended habitat for euryhaline estuarine and marine species, a function particularly important during low-flow and drought periods when the REI breaks down and / or the estuary may be cut off to the sea. Weirs, impoundments and other instream obstacles in the transition zone may greatly reduce the availability and benefits of this habitat to estuarine-associated species. This includes a reduction in the transport of production and detritus from the transition zone to the estuary downstream. Recruitment of estuarine-associated species will be limited to *G. aestuaria* and similar animals that are able to complete their entire lifecycle in this habitat. Alien and extralimital fish may have a similar impact. Many of these fish are predatory, outcompete their indigenous counterparts and thrive in transitional waters where they can provide an effective barrier to any larval or juvenile fish and invertebrates trying to recruit from downstream. Identifying these freshwater—estuarine transitions zones requires that the relevant lead agents i.e. Department of Water and Sanitation and Department of Environmental Affairs collaborate more closely on issues within these support areas that can potentially impact on estuaries.

## Conclusion

A total amount of 4 698 823.6 ha (3.9% of South Africa) of inland aquatic ecosystems, including inland wetlands, estuaries and some river channels in the National Wetland Map version 5, as well as an artificial wetlands data layer, have been mapped as part of the South African Inventory of Inland Aquatic Ecosystems (SAIIAE). The datasets and associated attributes have informed the National Biodiversity Assessment for 2018, as well as the Sustainable Development Goal reporting for indicator 6 through the Department of Water and Sanitation to the United Nations Environmental Programme. Significant effort is required to improve the confidence of the representation of the inland wetlands in the future updates of the National Wetland Map.

## Acknowledgements

Data sources have been duly acknowledged in the South African Inventory of Inland Aquatic Ecosystems (SAIIAE) Report (Van Deventer et al., 2018a) and journal paper (Van Deventer et al., 2018b). This study was funded by the Parliamentary Grant funding of the CSIR (Project EEEO053), funding allocated by the South African National Biodiversity Institute (SANBI) to the National Biodiversity Assessment for 2018 (NBA 2018) and the Water Research Commission (WRC) under the project WRC K5/2546 ‘Enabling more responsive policy and decision making in relation to wetlands through improving the quality of spatial wetland data in South Africa’, Desktop Provisional EcoClassification of the Temperate Estuaries of South Africa. WRC Report No. K5/2187, and A multi-sectoral Resource Planning Platform for South Africa’s estuaries. Water Research Commission Report No K5/2464.

ICLEI collaborated strongly with SANBI and the CSIR, and funded the mapping and review of the Eden District Municipality in the Western Cape. The mapping of wetlands in the Amathole (EC), Cape Winelands (WC), Ehlanzeni (MP) and Umgungundlovu (KNZ) District Municipalities had been funded by the Global Economic Fund 5 (GEF 5). The National Research Foundation (NRF) funded a number of internships to the CSIR. The WRC Project K5/2545 titled ‘Establishing remote sensing toolkits for monitoring freshwater ecosystems under global change’ contributed the updated wetlands data for Hogsback and Tevredenpan study areas.

A number of data capturers have spent months on improving the representation of inland aquatic ecosystem types for districts and provinces. We acknowledge the hard and tedious work of the following people: Ms Millicent Ketelo Dinala (SANBI), Ms Ridhwannah Gangat (CSIR, SANBI), Ms Kedibone Lamula (SANBI), Mr Leolin Qegu (NRF, CSIR & SANBI), Mr Phumlani Zwane (CSIR), Mr Mthobisi Wanda (ICLEI), Mr Gcobani Nzonda (SAEON), Mr Frikan Erwee (SANBI), Ms Carla-Louise Ramjukadh (CSIR), Mr Nhlanganiso Biyela (SANBI), Mr John April (NRF-CSIR), Ms Bongiwe Simka (NRF-CSIR), Ms Sinekhaya Maliwa (NRF-CSIR), Dr Heidi van Deventer (CSIR); Mr Tumisho Ngobela, Ms Kate Snaddon and Mr Dean Ollis from the Freshwater Consultancy Group (FCG) for the integration and improvement of the Cape Winelands District; interns from the Department of Water and Sanitation Directorate: Spatial & Land Information Management (SLIM) under supervision of Ms Carey Rajah for mapping systems in the North-West Province. Dr Mervyn Lötter and Mr Hannes Marais from the Mpumalanga Tourism and Parks Agency (MTPA) are particularly thanked for pulling the wetlands of the Mpumalanga Province together and fixing the data at a desktop level during their review of the province.

We are indebted to other reviews of the inland wetlands mapped, including Dr Brian Colloty (Scherman, Colloty and Associates), Ms Nancy Job (SANBI), Ms Kate Snaddon (FCG) and Dr Nacelle Collins (FS DESTEA). Dr Andrew Skowno has integrated the artificial wetlands and contributed to the improvement and edge-matching of the provinces, whereas Dr Heidi van Deventer corrected the large dams and smaller errors thereafter.

Mr John April has mapped the transitional rivers as listed in Appendix I. Drs Taryn Riddin and Meredith Fernandes, as well as Ms Carla-Louise Ramjukadh, Dr Lara van Niekerk, Ms Fiona McKay and Prof Janine Adams spent hours mapping, reviewing and correcting the estuarine ecosystem types and extent. The KZN Department of Economic Development, Tourism and Environmental Affairs and The Oceanographic Research Institute funded mapping of the EFZs of KZN. Dr Stephen Lamberth, Dr Lara van Niekerk and Dr Steven Weerts compiled the list and extent of the transitional zones which Dr Heidi van Deventer implemented in NWM5. The inland wetlands reference committee has provided guidance through the update of the NWM, including Dr Nacelle Collins (Department of Economic, Small Business Development, Tourism and Environmental Affairs), Mr Dean Ollis (Freshwater Consultancy Group), Ms Nancy Job (SANBI), Dr Mervyn Lötter (Mpumalanga Tourism and Parks Agency), Dr Erwin Sieben (University of KwaZulu-Natal) and Ms Kate Snaddon (Freshwater Consultancy Group). Prof Moses Cho (CSIR) for sharing the LAI dataset and advising on appropriate use.

Lastly, we are grateful to the reviewers who provided comments and suggestions to the improvement of this manuscript.

## APPENDIX I: SUPPLEMENTARY XLSX SPREADSHEET WITH EXTENT OF WETLANDS PER PROVINCE, DISTRICT AND MUNICIPALITY

## APPENDIX II: TRANSITION RIVER LISTS

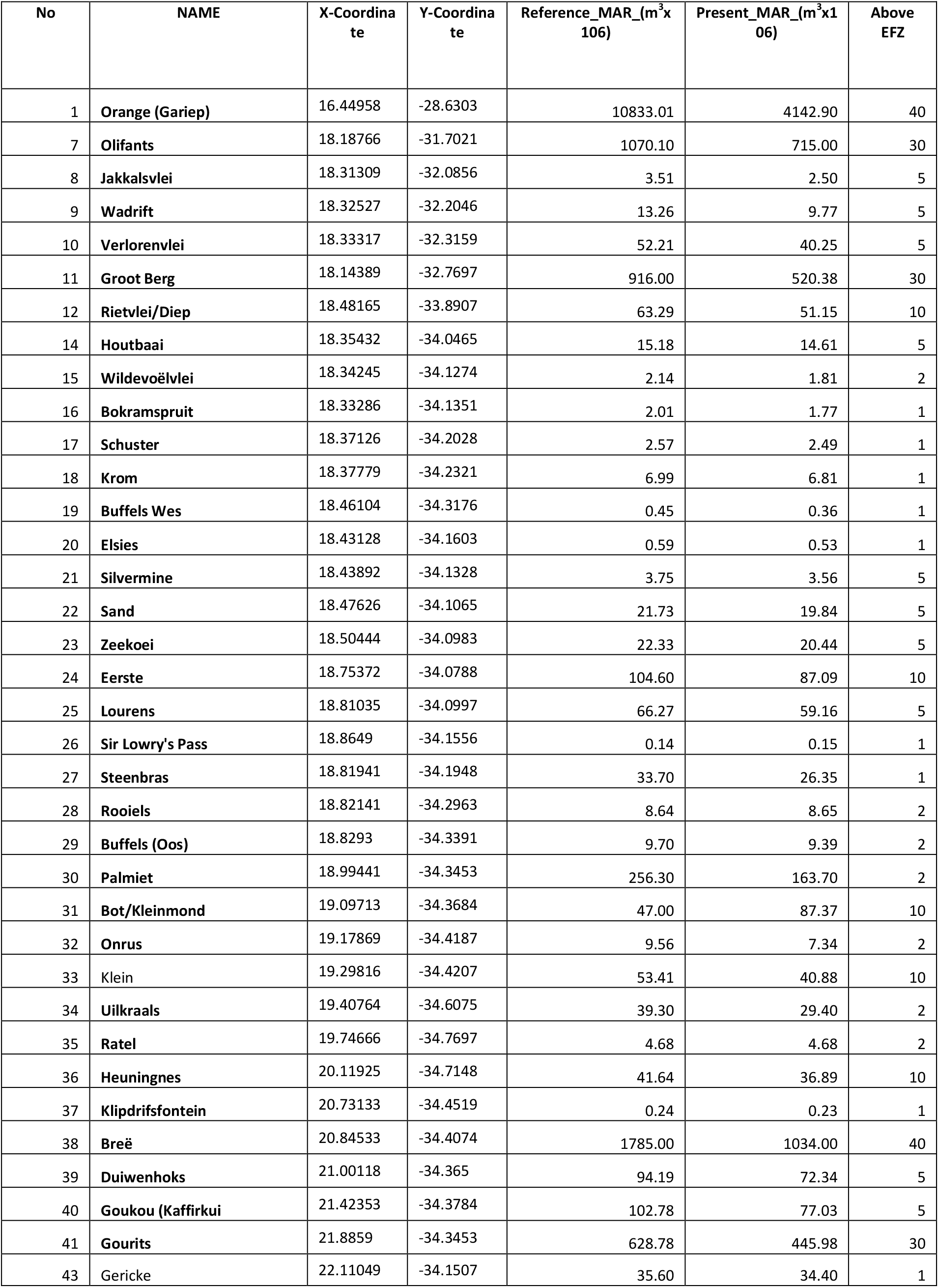
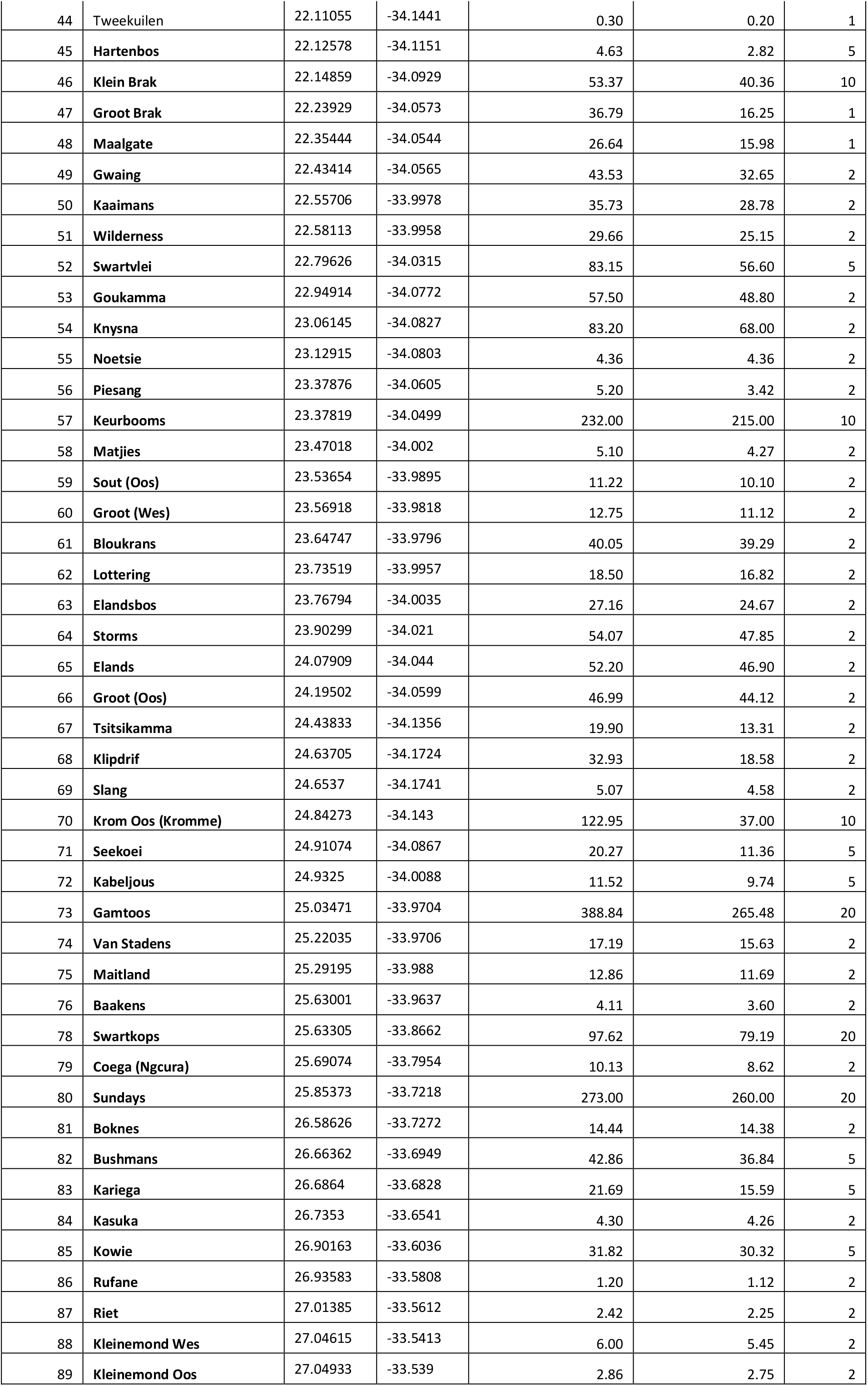
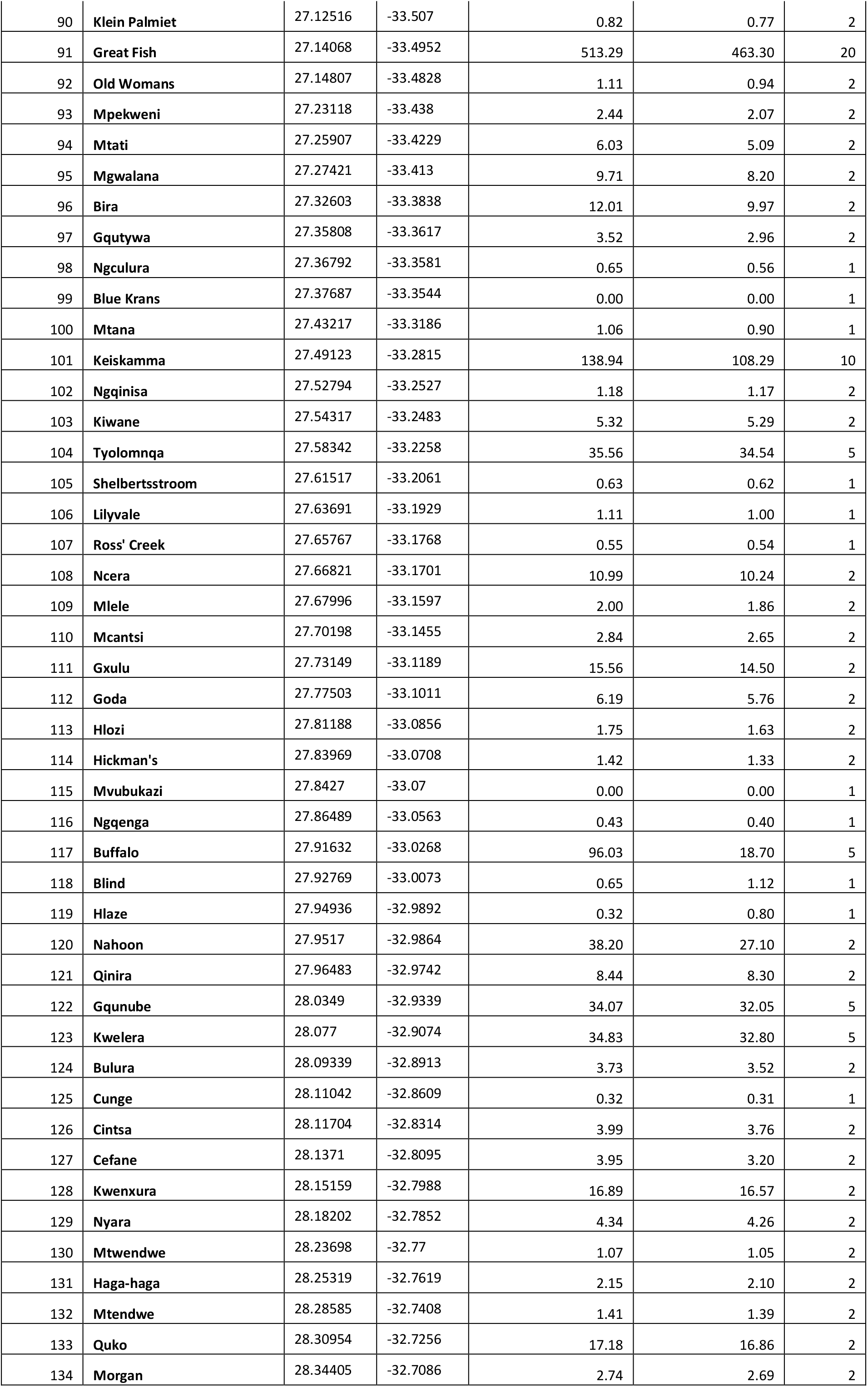
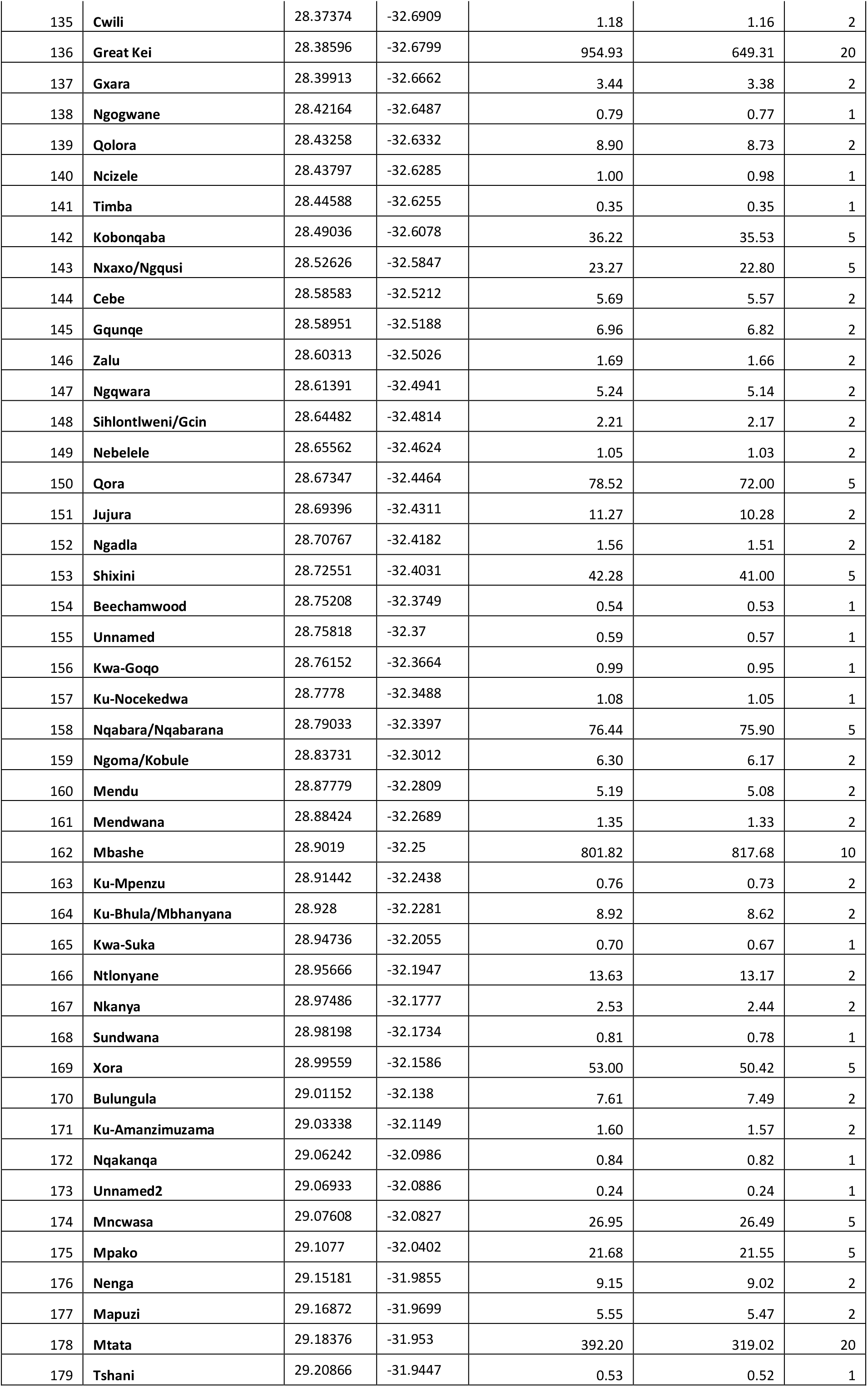
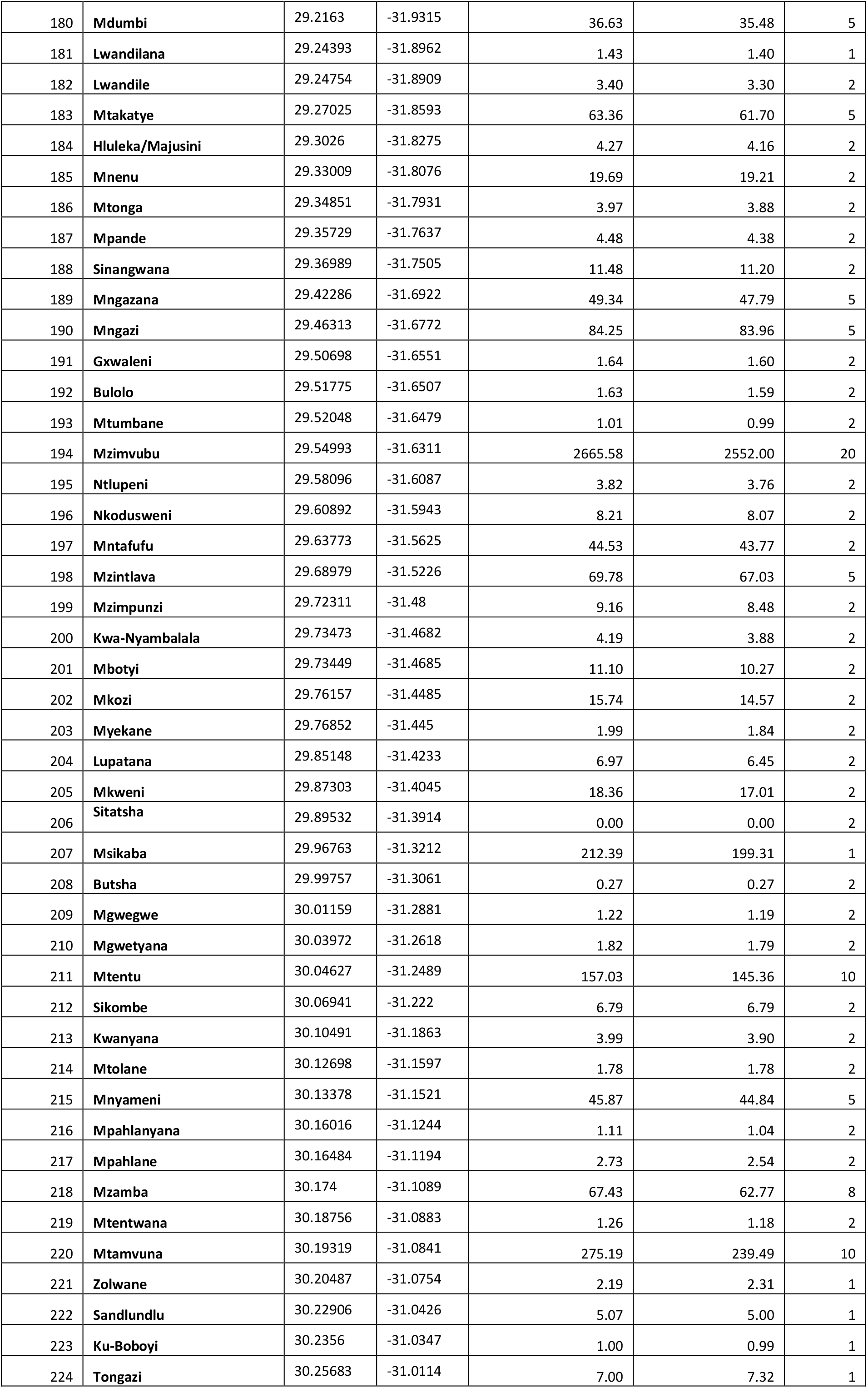
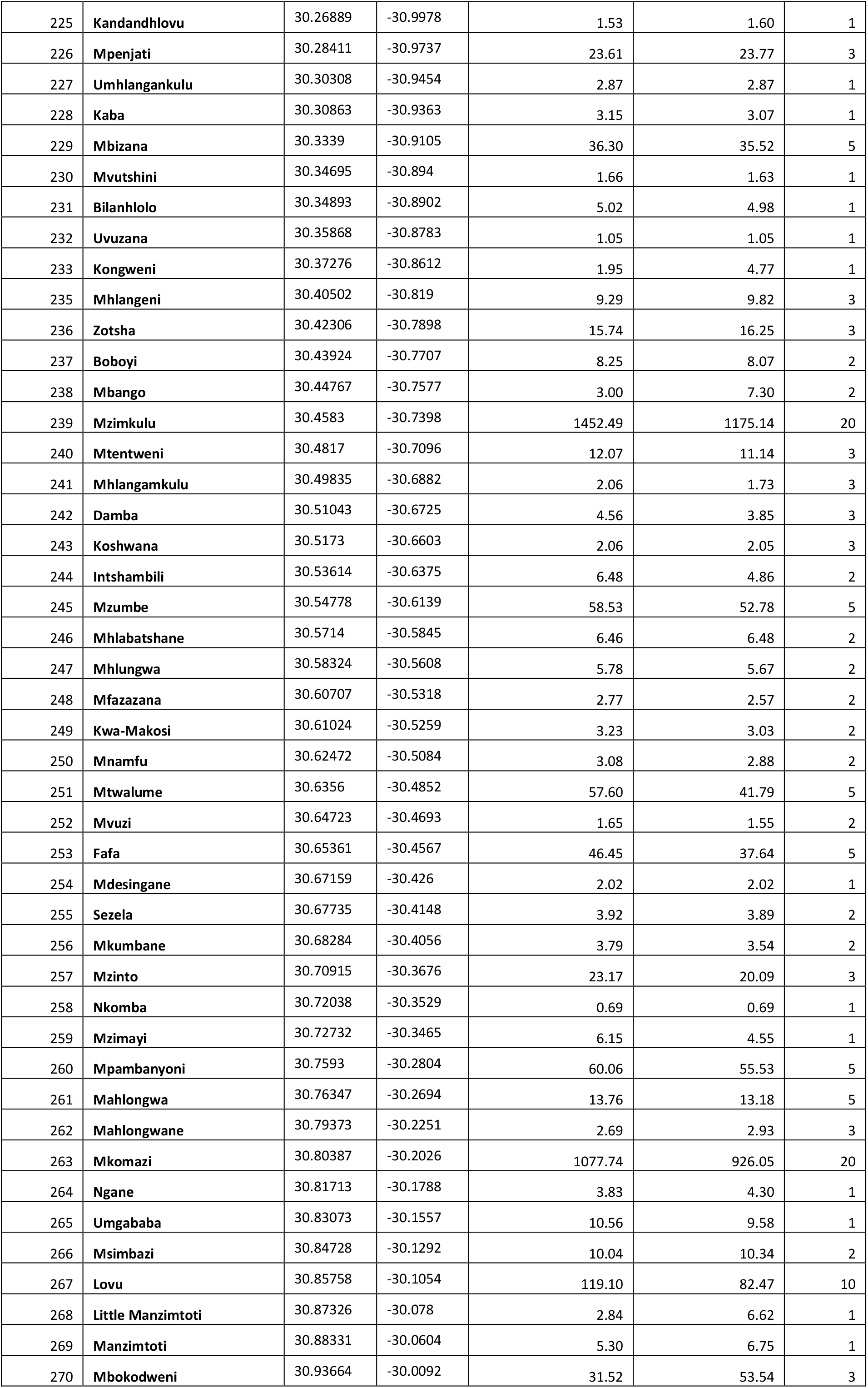
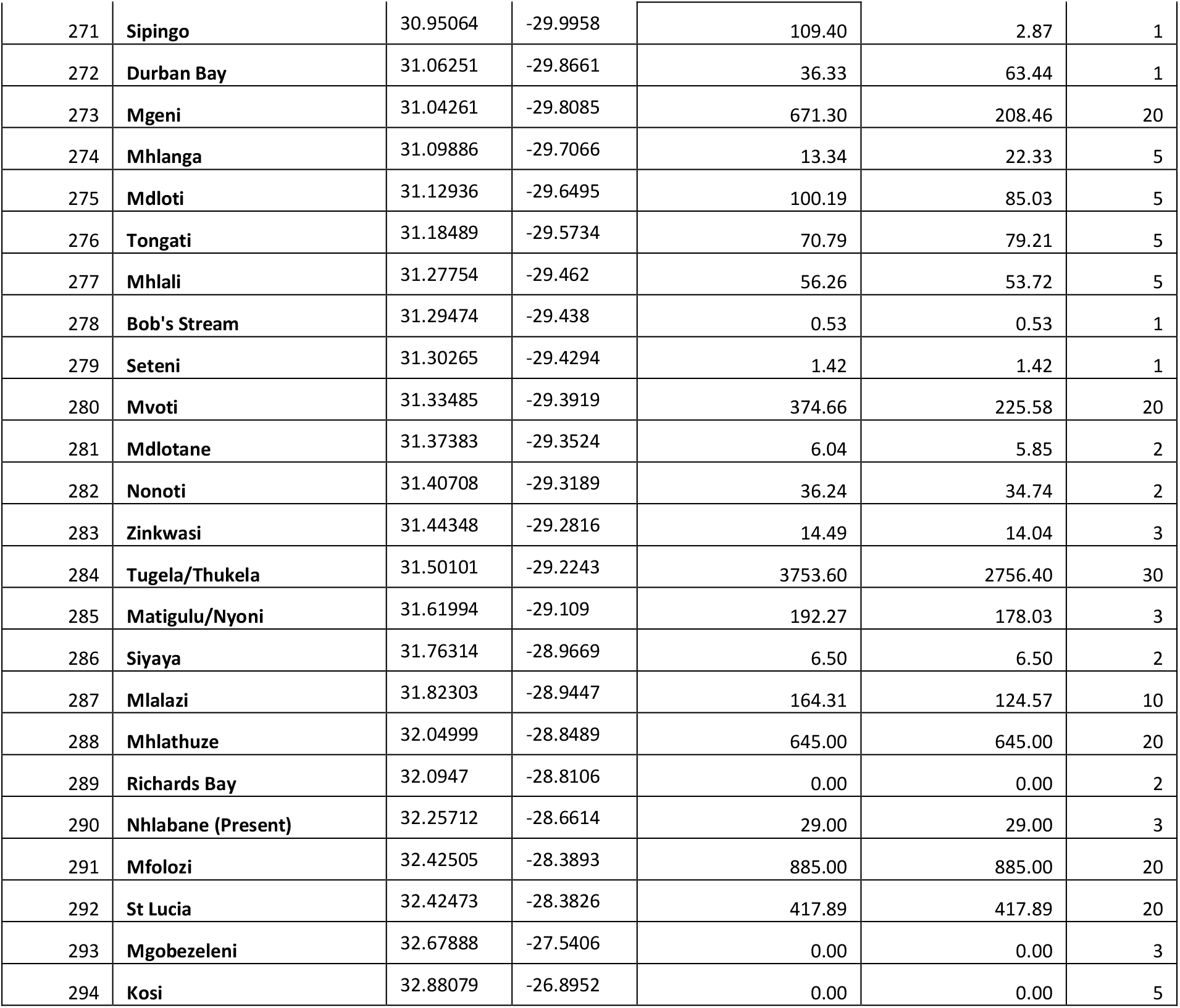

## Footnotes

1 *The extent of South Africa has been calculated as 121 973 563.7 ha using the provincial boundaries of the Municipal Demarcation Board (MDB) of 2011, however the marine reserves have been excluded from the shapefile by the CSIR.*

2 *Appendix I provides a full spreadsheet of the extent of wetlands per province, district and municipality, for use by Interested and Affected Parties as required.*

